# Polycomb contraction differentially regulates terminal human hematopoietic differentiation programs

**DOI:** 10.1101/647438

**Authors:** A. Lorzadeh, C. Hammond, F. Wang, D.J.H.F. Knapp, J.CH. Wong, J.Y.A Zhu, Q. Cao, A. Heravi-Moussavi, A. Carles, M. Wong, Z. Sharafian, M. Moksa, M. Bilenky, P.M. Lavoie, C.J. Eaves, M. Hirst

## Abstract

Lifelong production of the many types of mature blood cells from less differentiated progenitors is a hierarchically ordered process that spans multiple cell divisions. The nature and timing of the molecular events required to integrate the environmental signals, transcription factor activity, epigenetic modifications, and changes in gene expression involved are thus complex and still poorly understood. We now show that the more primitive types of human cells in this system display a unique repressive H3K27me3 signature that is retained by mature lymphoid cells but is lost in terminally differentiated monocytes and erythroblasts. Additional intervention data implicate that control of this chromatin state change is a requisite part of the process, whereby normal human hematopoietic progenitor cells make lymphoid and myeloid fate decisions.

## Main

Epigenetic modifications govern local chromatin activity and support activation or silencing of gene transcription through the regulation of chromatin structure and DNA accessibility^1,2^. Numerous cell differentiation processes are known to be accompanied by obligatory changes in chromatin structure mediated by proteins that specifically modify histones and thereby establish and maintain defined transcriptional regulatory states^3–7^. However, little is known about the details of epigenomic programing changes that promote or determine lineage restriction in normal tissues, or how such changes may contribute to the initiation and completion of terminal differentiation programs.

Hematopoiesis refers to the general process by which the different types of short-lived, mature blood cells are produced throughout life^8^. The accessibility of the cells directly involved has made this system an attractive one for identifying the types of epigenomic changes that characterize this process, the sequence in which they occur and those that are necessary^9–12^. At birth, human cord blood (CB) has been particularly useful for such studies, because it contains a self-maintaining population of hematopoietic stem cells (HSCs) that regenerate the entire system in transplanted receptive hosts lifelong as well as a full spectrum of derivative progenitors with reduced proliferative capacity and different lineage options. These progenitor cells then undergo further restriction of their proliferative and lineage potentials to eventually allow the initiation of a single lineage program and the production of one of the many different types of mature blood cells^13,14^. Previous studies have suggested that the process of lineage restriction involves the epigenetic demarcation of specific genomic regions that make them accessible to particular transcription factors to ultimately activate single lineage-specific gene expression programs^10,15^. However, neither the molecular details of these steps nor their dynamics in relation to cell phenotype changes have been delineated.

Previous studies using chromatin immunoprecipitation sequencing (ChIP-seq) of sites of permissive histone modifications and transposase-accessible chromatin sequencing (ATAC-seq) have provided insight into the identity of regulatory regions in the genomes of various phenotypically defined primitive subsets of mouse and human hematopoietic cells that change during their differentiation^9–11^. Together, these findings have suggested a model in which certain enhancers are initially “primed” in primitive multi-potent hematopoietic cells by an acquisition of “permissive” histone 3 lysine 4 mono-methylation (H3K4me1) modifications, followed by a gain of additional histone 3 lysine 27 acetylation (H3K27ac) modifications by the same histones to enable terminal differentiation programs to become activated^9^. However, the potential existence of “stage-specific” as well as lineage-specific changes in the chromatin as an integral part of the process ^14,16,17^, has remained uncharacterized, particularly in human hematopoiesis.

To obtain such information, we isolated 8 previously well characterized, phenotypically defined subsets of normal human CB cells at high purity^14,16,17^ and generated detailed genome-wide datasets for permissive and repressive histone modifications, sites of DNA methylation and transcriptomes from them. Four of these were highly enriched in cells with progenitor activity but different lineage output capabilities^16^ and together constitute the bulk of all of the CD34+ CB cells. They consisted of a CD38-subset, which contains the most primitive cells and all HSCs, and 3 distinct subsets within the CD38+ subpopulation of CD34+ cells, the so-called common myeloid progenitors (CMPs), granulocyte-macrophage progenitors (GMPs) and megakaryocyte-erythroid progenitors (MEPs). The 4 “mature” cell types analyzed were low-density, circulating monocytes, erythroblasts, B cells and T cells.

Analysis of repressive chromatin states in the 4 (CD34+) progenitor populations showed these all shared a nearly identical H3K27me3 signature (gene promoter Spearman correlation R>0.97). This signature included large organized chromatin K27-modification domains (LOCKs) that were also present in the B and T cell isolates, but absent from the monocytes and erythroblasts. The LOCKs present in all of the progenitor fractions examined as well as in the mature lymphoid cells were also found to be co-marked with H3K9me3 and located within lamin-associated domains. The CpGs within the B and T cells also resembled the CD34+ progenitor fractions in their general hypermethylation status in comparison to the CpGs of the co-isolated monocytes and erythroblasts, consistent with the acquisition of an altered heterochromatic state by the latter two cell types. Analysis of the enhancer landscape of these 8 populations revealed that a majority of traditionally defined active enhancers found in differentiated cell types were already evident within their progenitors at a hypermethylated state, and those unique to terminally differentiated cells, were found almost exclusively within the boundaries of super-enhancers. From these findings, we propose a model in which a genome-wide contraction of heterochromatin is a critical step in the process by which human hematopoietic progenitor cells with lympho-myeloid potential lose their lymphoid potential.

### Identification of a chromatin signature shared by progenitors and mature lymphoid cells but not mature myeloid cells

We first undertook a comprehensive mapping of the epigenetic and transcriptional states of historically defined immunophenotypes of human CB cells using our low input ChIP-seq protocol^18,19^ to identify H3K4me3, H3K4me1, H3K27me3, H3K27ac, H3K36me3, and H3K9me3 sites genome-wide, plus whole genome bisulfite sequencing and RNA-seq protocols following IHEC standards^20^ (**Fig. 1A** and **S1A**). A standardized analytical pipeline was then applied to qualify and analyze the resulting data (see methods).

**Figure 1.**
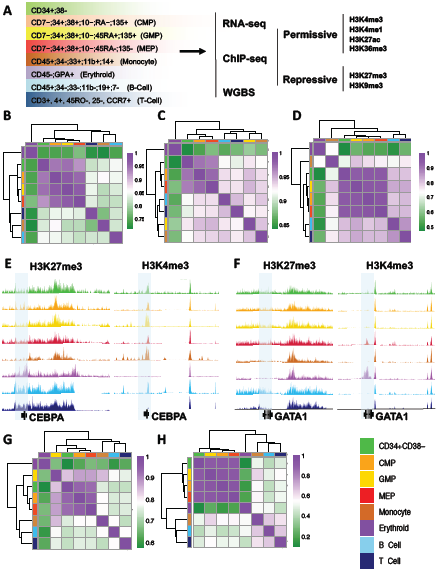
H3K27me3 signatures are shared across functionally distinct human CD34+ hematopoietic progenitor populations. **A)** Schematic of the experimental design and colour legend (**shown in the bottom right corner**) for the 8 cell types analyzed. Unsupervised hierarchical clustering and heatmap of pairwise Spearman correlations for protein coding gene RPKM values **(B)**, H3K4me3 coding gene promoter (± 2 Kb) density **(C)**, and H3K27me3 coding gene promoter density **(D)** across cell types indicated by colour. Genome browser view of H3K27me3 (left panel) and H3K4me3 (right panel) density tracks across cell populations indicated by track colour at the *CEBPA* **(E)** and *GATA1* **(F)** locus. Unsupervised hierarchical clustering and heatmap showing pairwise Spearman correlation of H3K4me3 **(G)** and H3K27me3 **(H)** densities genome-wide.

RNA expression profiles for each of the progenitor populations analyzed (CD34+CD38-cells, CMPs, GMPs and MEPs) were more highly correlated with one another (Spearman R >0.92) than with any of the 4 more mature CB cell types examined; i.e., monocytes, erythroblasts, and B and T cells (average Spearman R = 0.76, 0.83, 0.84 and 0.84, respectively, **Fig. 1B** and **S1B**). These confirm relationships also evident in previously published datasets^3^. Expression signatures derived for each progenitor subset were also in agreement with previously published features of these same subsets (methods and see **Fig. S2A** and **B**). For example, pathway and gene enrichment analysis of uniquely upregulated transcripts in GMPs showed these were enriched in terms related to leukocyte differentiation, inflammation response and regulation of immune response (Benjamini *q*-value <10e^-20^, **Fig. S2C**), and included the CD135 cell surface marker. Transcripts uniquely upregulated in MEPs were enriched in terms related to myeloid lineage differentiation (Benjamini *q*-value <10e-3) (**Fig. S2D**). Genes up-regulated in CD34+CD38-cells and CMPs as compared to the 4 populations of more differentiated cells examined were enriched in terms related to hematopoiesis regulation and differentiation (Benjamini *q*-value <10e^-3^) (**Fig. S2E** and **F**).

Examination of the chromatin state of the 8 different cell types examined here showed H3K4me3 and H3K36me3 densities correlated with expected transcript levels and cell type-specific signatures (**Fig. 1C, E-G, S2G** and **H**). H3K4me3 densities at promoters of genes that were expressed at different levels in each of the different progenitor subsets, or between them and the 4 later cell types, also showed expected relationships (**Fig. S3A**). In contrast, H3K27me3 occupancy was nearly identical across all of the 4 different progenitor subsets (**Fig. 1D** and **1H**; promoter Spearman R >0.97), and cell-type specific signatures were apparent only in the mature cell types (**Fig. 1E** and **F**). However, the H3K27me3 densities at promoters in the progenitor fractions were more significantly correlated with those in the lymphoid cells as compared to either the monocytes or the erythroblasts (**Fig. 1D**). In fact, RNA expression, H3K4me3 and H3K27me3 signatures at promoters in the erythroid precursor population all showed the lowest correlation (average Spearman R <0.56) with the corresponding features in all 4 progenitor populations, including the MEPs (**Fig. 1D**). Moreover, a comparison of the H3K27me3 densities within the promoters of genes that were differentially expressed in GMPs and MEPs showed these were not significantly different (**Fig. S3B**). In contrast, genes whose expression appeared downregulated in monocytes and erythroblasts compared to GMPs and MEPs, respectively, showed a significant gain of H3K27me3 density at the corresponding promoters (2-sided t-test, *p* <2.2 × 10^−16^).

### Terminally differentiated monocytes and erythroblasts exhibit a genome-wide contraction of H3K27me3 density

Examination of the patterns of H3K27me3 occupancy in the chromatin of the 4 different types of mature blood cells analyzed (**Fig. 1D** and **1H**) showed both the monocytes and erythroblasts contained a significantly reduced overall frequency of H3 histones that were methylated at K27 (30-52%) (**Fig. S3C**). This was explained in part by the observation of a pronounced contraction of the extensive contiguous stretches of chromatin (domains) containing the H3K27me3 mark characteristic of the 4 progenitor populations. The remaining H3K27me3 in the monocytes and erythroblasts displayed a more punctate structure reminiscent of that seen in pluripotent cell types^21^ (**Fig. 2A-E**). Immunoprecipitated (IP) fragment distributions at promoters^19^ (± 2 Kb of transcription start sites (TSS)) were also significantly different in the monocytes and erythroblasts in comparison to the 4 progenitor subsets or to either the mature B or T cells present in the same samples (kolmogorov–smirnov test, *p*<7×10^−12^; **Fig. 2C**). Examination of H3K27me3 promoter distributions in all of these cell phenotypes also revealed an increase in the proportion of H3K27me3-marked histones at promoters in the monocytes and erythroblasts compared to the progenitors from which they are thought to be most immediately derived (GMPs, **Fig. S3D;** and MEPs, **Fig**.**S3F**), without a measurable change in H3K4me3 promoter occupancy (**Fig. S3E** and **G**).

**Figure 2.**
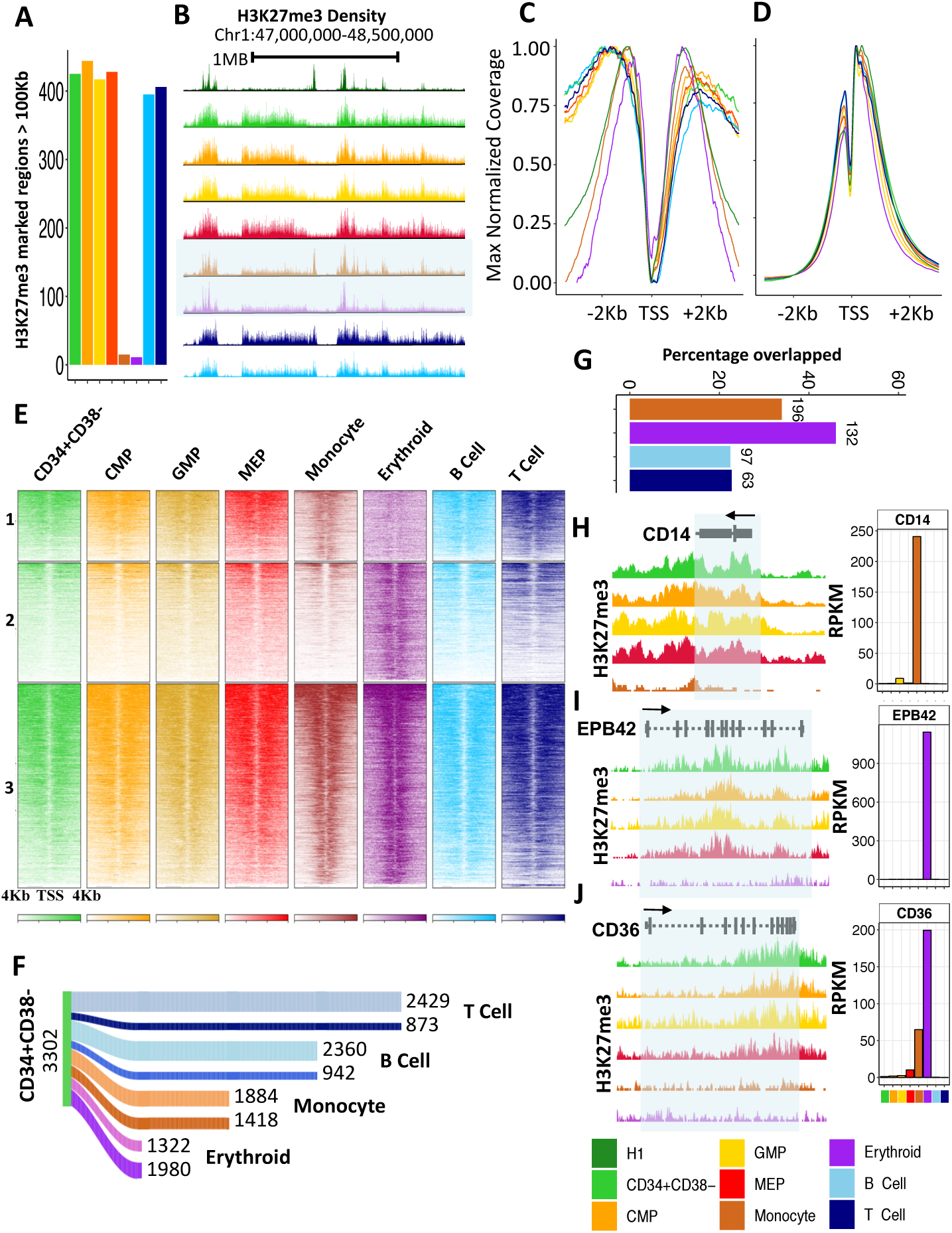
Myeloid differentiation is associated with genome-wide contraction of H3K27me3 –marked domains. **A)** Number of H3K27me3-marked genomic regions with widths >100 Kb across cell types indicated by the colour legend on the bottom right. **B)** Genome browser view of H3K27me3 density profiles over a 2 Mb window of chromosome 1 highlighting contraction of H3K27me3 in the monocyte (brown) and erythrblast (purple) cells. Maximum value normalized H3K27me3 **(C)** and H3K4me3 **(D)** density at coding gene promoters (± 2 Kb of TSS). **E)** Heatmap of H3K27me3 densities coloured by cell type at gene promoters and flanks (TSS ± 4 Kb) for regions marked by H3K27me3 in monocytes but not erythroblasts (**panel 1**), in erythroblast but not monocytes (**panel 2**), or in both (**panel 3**). Shading indicates increased density. **F)** Sankey diagram of number of genes retaining or losing H3K27me3 at their gene body in monocytes, erythroblasts, B and T cells compared to the CD34+CD38-population. (Dark shade: loss of H3K27me3, light shade retention of H3K27me3). **G)** Percentage of overlap of genes that lost H3K27me3 within up-regulated gene bodies in monocytes, erythroblasts, B and T cells compared to CD34+CD38-cells (the absolute number of overlapping genes is shown on top of each bar). Genome browser view of the H3K27me3 density at *CD14* **(H)**, *EPB42* **(I)** and *CD36* **(J)** locus. Expression (RPKM) of each gene across all profiled cell types are shown in the right panel.

To examine the functional consequence of the global alteration in H3K27me3 occupancy, we examined the levels of H3K27me3 within gene bodies in relation to transcript levels in all 8 cell types (**Fig. 2F**). Consistent with the genome-wide patterns, the monocytes and erythroblasts showed a loss in H3K27me3 density on more genes (1,418 and 1,980, respectively) than the B and T cells (942 and 873, respectively) when both sets were compared to the most primitive CD34^+^CD38^-^ progenitor compartment (**Fig. 2F**). In addition, loss of H3K27me3 correlated with a greater proportion of upregulated genes in monocytes and erythroblasts compared to the B and T cells (**Fig. 2G**). Among the genes that lost H3K27me3 and were upregulated were well-known markers of monocyte and erythroid lineage differentiation including CD14, EPB42 and CD36 (**Fig. 2H-J**).

In addition to the differential gene-specific losses of H3K27me3 in the monocytes and erythroblasts, there was also an associated loss of many large organized chromatin K27me3 domains (LOCKs)^22^ (**Fig. 3A** and **B**) that were, nevertheless generally retained in the B and T cells (**Fig. 3A** and **B**). A majority (average 87%) of the few remaining LOCKs identified in the monocytes and erythroblasts overlapped with LOCKs present in the 4 progenitor populations (**Fig. 3B**). Even within the LOCKs retained in the monocytes and erythroblasts, H3K27me3-marked regions showed evidence of contraction, with an average width of 52 Kb compared to 268 Kb seen in related progenitor subsets (GMPS and MEPs, **Fig. 3C-E**).

**Figure 3.**
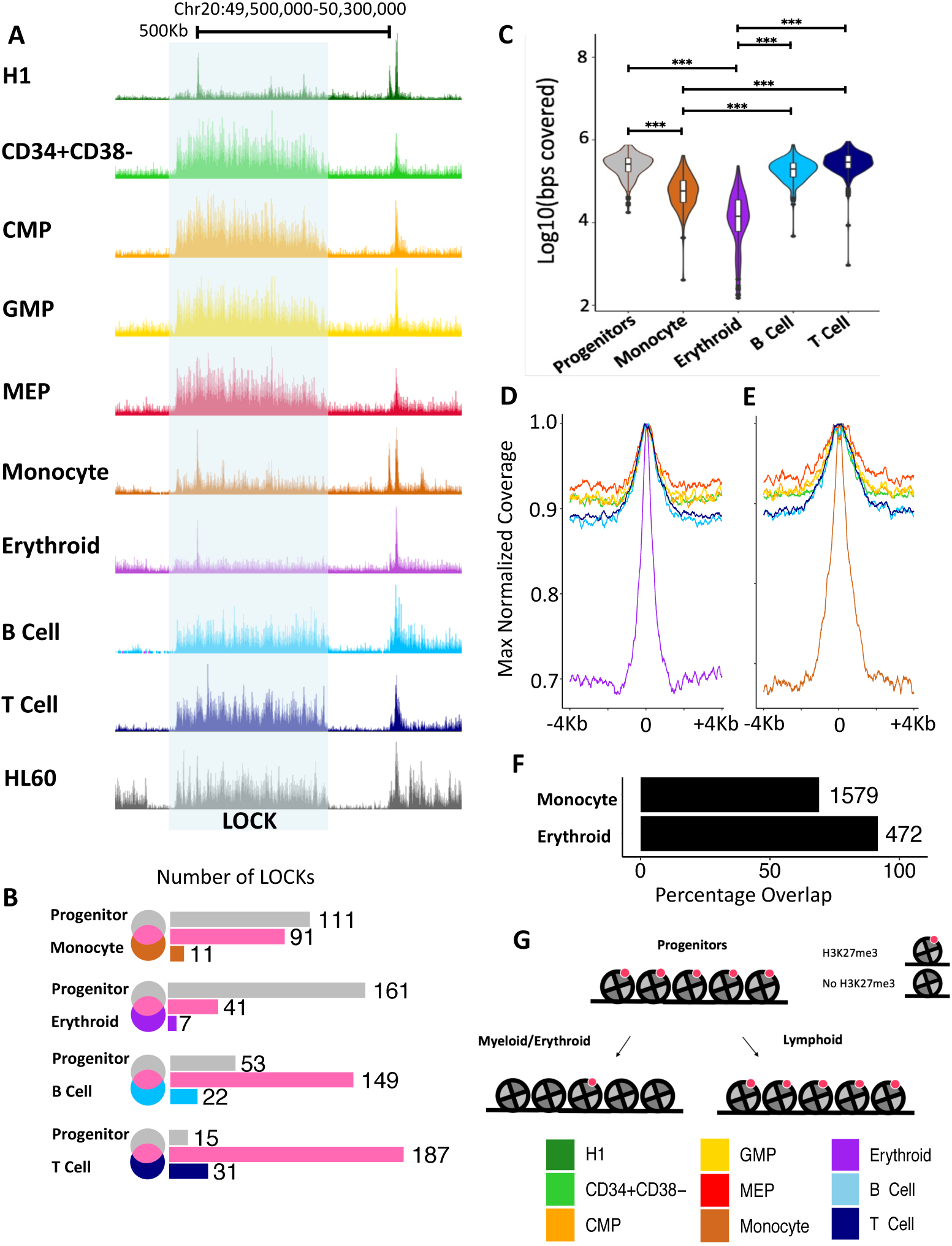
Contraction of H3K27me3 domains seen in monocytes and erythroblasts coincides with a loss of LOCKs in progenitors. **A)** Genome browser view of H3K27me3 density on chromosome 20 across cell types as indicated by the colour legend (bottom right). The shaded box indicates H3K27me3 LOCK (FDR <0.05) in progenitor cells that is absent in monocytes and erythroblasts. **B)** Venn diagram of LOCKs in all progenitors (CD34+CD38-; MEP; CMP; GMP) compared to those in monocytes, erythroblasts, B cells, and T cells. **C)** Violin plot of base pairs marked by H3K27me3 within LOCKs across cell types, as indicated (*** *p* <0.001). Maximum values for normalized H3K27me3 densities enriched within LOCKs in erythroblasts **(D)** and monocytes **(E). F)** Barplot showing the percentage overlap of H3K27me3 enriched regions within progenitors’ LOCKs also in monocytes and erythroblasts with H3K27me3-marked regions identified in H1 ESCs. **G)** Cartoon of contraction of H3K27me3-marked nucleosomes (red) during monocyte/erythroid differentiation. (*** *p* < 0.001)

The corresponding punctate signature observed in monocyte and erythroblasts is reminiscent of a chromatin state initially annotated in human embryonic stem cells (ESCs) that is lost following their differentiation^21^. This reversion of differentiated monocytes and erythroblasts to a chromatin state previously noted in undifferentiated ESCs was thus unexpected and prompted us to quantify the similarities between the LOCKs present in all the CB cell types analyzed and ESCs. This showed a majority of H3K27me3 marked regions within the LOCKs in the progenitor subsets that were also present in the monocytes and erythroblasts were also marked by H3K27me3 in the ESCs (**Fig. 3F**). These data thus provide evidence of a genome-wide contraction of H3K27me3 density during the process by which certain CB progenitors generate mature progeny (**Fig. 3G**) that appears to be lacking in the lymphoid restriction process.

### H3K27me3 LOCKs are enriched in lamina-associated domains (LADs)

To further characterize the H3K27me3 LOCKs lost during the terminal differentiation of monocytes and erythroblasts, we next examined the co-occurrence of other histone modifications and DNA methylation in these same regions. For this, we applied ChromHMM^23^ to generate an 18-state model based on H3K4me1, H3K4me3, H3K27me3, H3K27ac, H3K36me3, H3K9me3 occupancy across all of the 8 CB cell types examined (**Fig. 4A**). H3K9me3 was the most highly enriched mark within H3K27me3-defined LOCKs in all 8 cell types (**Fig. 4B** and **C**). Furthermore, LOCKs with or without H3K9me3 were found to be enriched in lamina associated domains (LADs) in both the progenitor subsets and in the B and T cells, but not in monocytes or erythroblasts (**Fig. 4A, D, E** and **S4A**). In contrast, regions enriched in H3K9me3 alone were strongly associated with LADs in all 8 cell types (**Fig. 4A** and **S4A**). A majority of LOCKs across all cell types overlapped with laminB binding sites (> 55%) and these appeared to be specifically lost in monocytes and erythroblasts (**Fig. 4F** and **4G)**. H3K9me3 occupancy, like that of H3K27me3, was significantly reduced in the monocytes and erythroblasts compared to the progenitor subsets (2-sided t-test *p*<2.2 × 10^−16^, **Fig. 4H**). H3K9me3 is also highly correlated between the 4 progenitor populations compared to any of the 4 mature cell types studies and there was a directional loss of H3K9me3 occupancy specifically in monocytes and erythroblasts (**Fig. S4B and C**).

**Figure 4.**
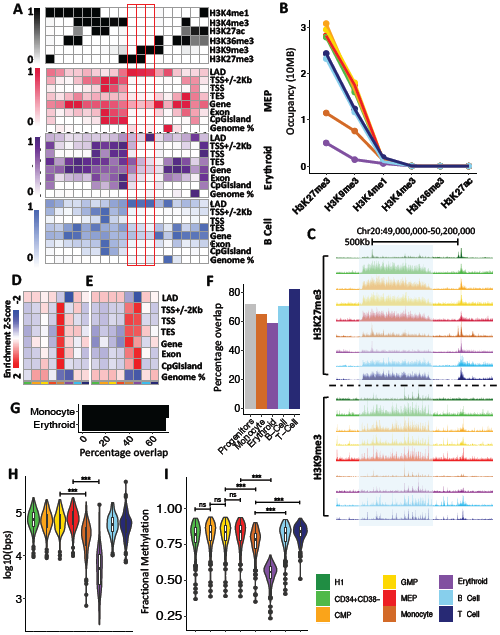
H3K27me3 LOCKs lost during myeloid differentiation are associated with LADs in CD34+ progenitor cells. **A)** Emission probabilities by histone mark for the 18 states of the ChromHMM chromatin state model across all cell types (upper panel). Enrichment of chromatin states within genomic features for MEPs (red), erythroblasts (purple) and B cells (blue). **B)** Cumulative number of bases marked by indicated histone modification within progenitor population LOCKs as indicated by the colour legend (bottom right). **C)** Genome browser view of H3K27me3 and H3K9me3 density on chromosome 20 across cell types indicated by the colour legend (bottom left). The shaded box indicates a H3K27me3 LOCK (FDR < 0.05) present in progenitor cells but lost in monocytes and erythroblasts. Heatmap of z-score enrichment of ChromHMM identified polycomb repressed **(D)** and dually repressed **(E)** chromatin states at genomic features. **F)** Barplot showing percentage of LOCKs overlapping with laminB binding site. **G)** Barplot showing percentage overlap of LOCKs lost in monocyte or erythroblasts with laminB binding site. **H)** Violin plot showing occupancy of H3K9me3 (log10(bps)) at LOCKs across all populations as indicated by the colour legend (bottom right). **I)** Violin plot of fractional CpG methylation levels within progenitors’ LOCKs across all populations as indicated by the colour legend (bottom right) (*** *p* < 0.001).

Consistent with the genome-wide reduction in heterochromatic states exhibited by the monocytes and erythroblasts, these two cell types showed reduced CpG methylation in their LOCKs compared to the 4 progenitor subsets and the B and T cells (2-sided t-test *p*<5.5 × 10^−8^; **Fig. 4I**). Erythroblasts particularly, but also the monocytes, showed reduced DNA methylation more broadly in chromatin states identified as polycomb-repressed and/or heterochromatin-repressed in the ChromHMM model (**Fig. S4D-G**). These findings are consistent with a previously reported reduction in DNA methylation of monocytes and neutrophils relative to different progenitor populations^12^. They also corroborate a previous relationship between DNA methylation and H3K9me3 densities^24–26^. In the present context, they also suggest that human CD34+ hematopoietic progenitors share a higher order chromatin structure that is associated strongly with LADs and is enriched in sites of H3K27me3 and H3K9me3.

### H3K27me3 contraction is associated with reduced *BMI1* expression

We next asked whether there were any similarities in the expression of polycomb group (PcG) proteins in monocytes and erythroblasts in comparison to ESCs. B cell-specific Moloney murine leukemia virus integration site 1 (*BMI1*), a component of the polycomb repressive complex 1 (PRC1), was the only PcG gene for which consistently lower transcript levels were seen in all 3 of these cell types compared to the other 6 types profiled here (**Fig 5A**), and in the mouse (**Fig 5B**)^9^. Transcriptional repression of *BMI1* has been previously associated with a loss of both H3K27me3 and H3K9me3, and a concomitant reduction in heterochromatin^27,28^. Likewise, inhibition of *BMI1* expression *in vivo* compromised hematopoietic progenitor self-renewal^29,30^. These observations suggest that the contraction of H3K27me3 observed in monocytes and erythroblasts and shared with ESCs, is associated with reduced expression of *BMI1*, a PRC1 complex member previously implicated in the maintenance of H3K27me3 marks.

**Figure 5.**
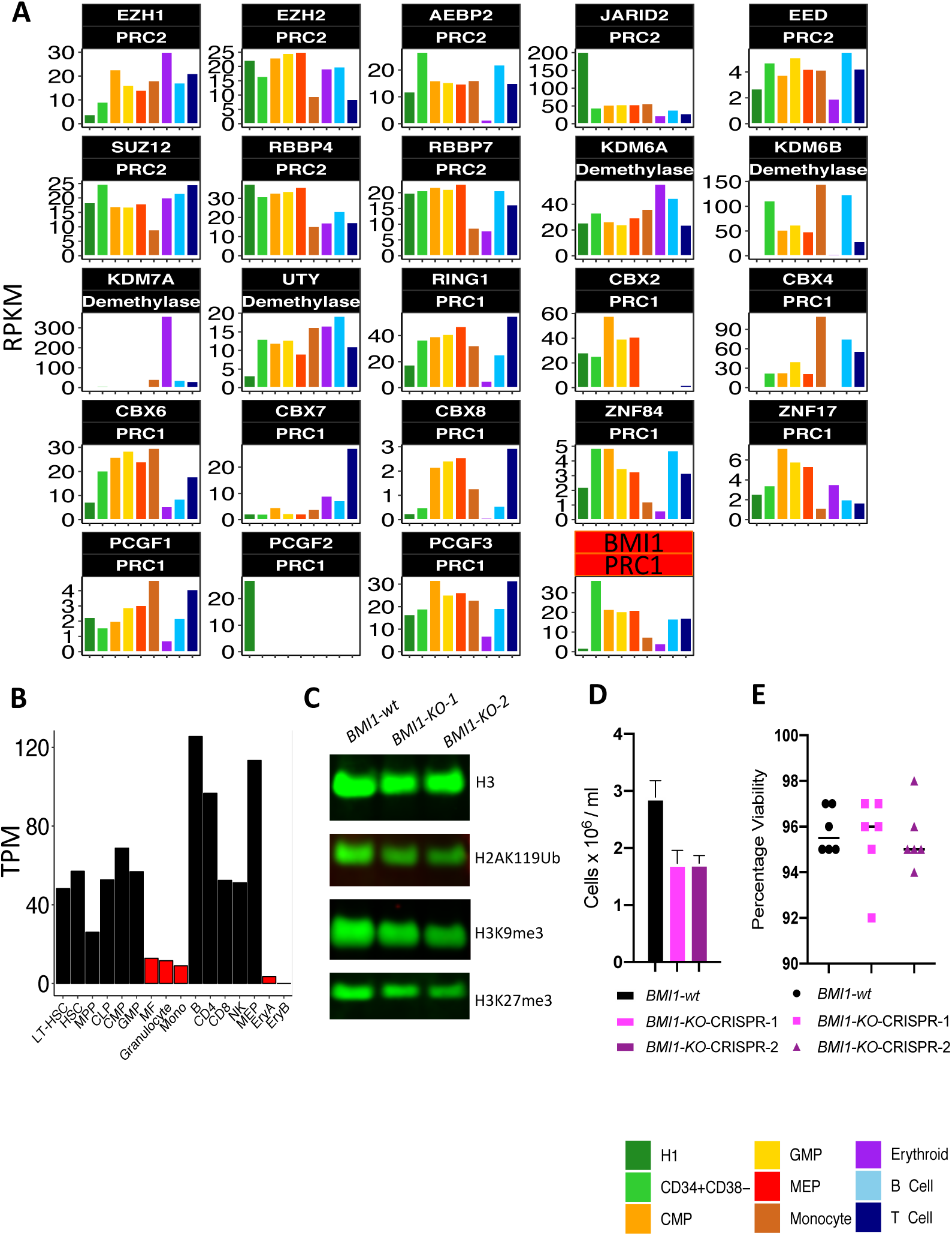
*BMI1* is expressed at a lower level in monocytes, erythroblasts and ESCs compared to other cell types. **A)** Expression (RPKM) of *PRC1, PRC2* and H3K27me3 demethylase across cell types as indicated by the colour legend (bottom right). **B)** Expression (TPM) of *Bmi1* in mouse hematopoietic cells. **C)** Global measurement of H3, H2AK119Ub, H3K9me3 and H3K27me3 by western blot across *BMI1-wild type* (*wt)* and *BMI1-knockout* (*KO*). **D)** Total number of cells at 3 days post CRISPR treatment across *BMI1-wt* and *KO*. **E)** Percentage of viable cells in **(D)**.

To investigate the role of *BMI1* expression on H3K27me3 maintenance, we first examined the ability of HL60 cells to maintain a progenitor-like H3K27me3 state at LOCKs (**Fig. 3A**) in the presence or absence of BMI1. We utilized CRISPR/cas9 to target the *BMI1* allele in two independent biological replicates and validated its knockout using a T7 endonuclease mismatch cleavage assay and at the protein level by western blot (**Table S1** and **Fig S5A-C**). As expected, given its role in mediating histone ubiquitination, *BMI1-KO* cells showed significant reduction in H2AK119 ubiquitination (H2AK119Ub) compared to control cells (**Fig 5C** and **S5B**). Loss of BMI1 was also accompanied by a decrease in global levels of both H3K27me3 and H3K9me3 and reduced proliferation (**Fig 5C-E** and **S5B, D** and **E**), confirming previous reports of the same effects^27,28^. ChIP-seq confirmed a genome-wide, rather than localized reduction of H3K27me3 in the *BMI1-KO* cells suggesting loss of H3K27me3 both within and outside of LOCKs (**Fig S5F-I**). These results confirm previously reported associations between *BMI1* loss and H3K27me3 reduction^27,28^.

### Hematopoietic progenitor subsets share lineage-specific enhancers marked by H3K27ac

We next sought to identify and compare the enhancer states of the 4 progenitor populations and the 4 more mature cell types examined. Accordingly, we identified the H3K27ac- and H3K4me1-marked regions in each population and used the results to create a catalogue of active (H3K27ac and H3K4me1) and primed (H3K4me1) enhancers. Primed enhancers were relatively consistent across all 4 progenitor populations (Spearman, average R >0.8; **Fig. 6A**) with active enhancers showing a higher degree of progenitor specificity (Spearman, average R >0.52; **Fig. 6B** and **S6C**). The progenitor populations also showed consistently a higher number of total enhancers, as measured by the sum of both the active and primed enhancer states, by comparison to the 4 more mature cell types (**Fig. S6A**). In contrast, the number of active enhancers was higher in 3 of the 4 more mature cell types (2-sided t-test, *p* =0.026), the exception being the erythroblasts that had the lowest number of active enhancers compared to all other cell types (**Fig. S6B**).

**Figure 6.**
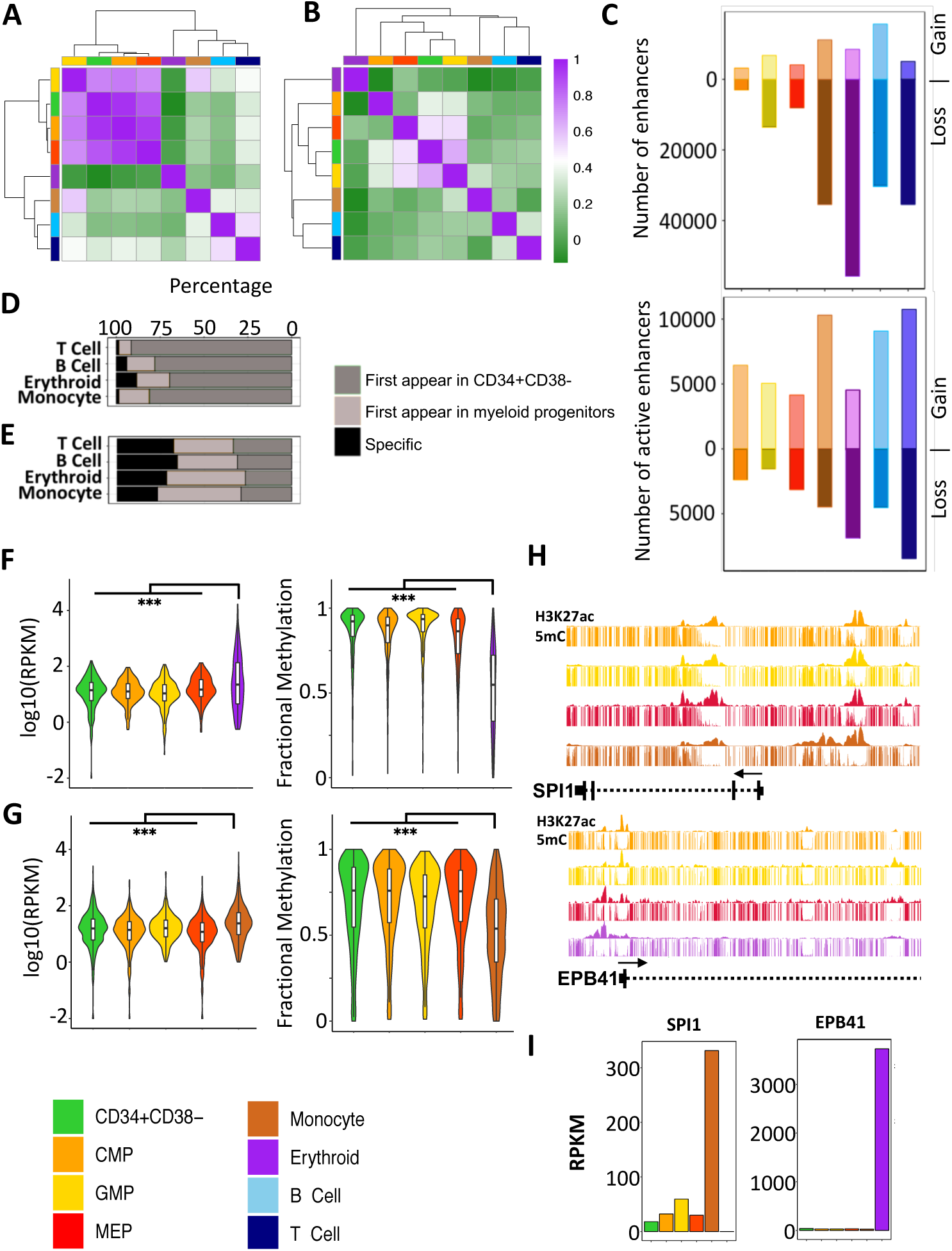
A majority of active enhancers in mature myeloid cells are marked by H3K27ac in CD34+ progenitor CB cell populations. Unsupervised hierarchical clustering and a heatmap of pairwise Spearman correlations of H3K4me1 **(A)** and H3K27ac **(B)** densities across cell types (colour legend shown at bottom left). **C)** Number of total enhancers (top panel) and active enhancers (bottom panel) gained or lost in each cell type compared to CD34+CD38-CB cells. Percentage of enhancers **(D)** and active enhancers **(E)** in differentiated cells also in CD34+CD38-cells, myeloid progenitor cells, or uniquely in monocytes, erythroblasts, or B or T cells. Expression of genes (left panel) with active enhancers and fractional DNA methylation at active enhancers in erythroblasts (**F**) and monocytes (**G**) also in their inferred parental progenitor populations. **H)** Genome browser view of H3K27ac density at *SPI1* (top panel) and *EPB41* (bottom panel) locus. **I)** Expression (RPKM) of *SPI1* (left panel) and *EPB4* (right panel). (*** *p* <0.001).

We next traced the gain or loss of H3K27ac and H3K4me1 from the most primitive CD34+CD38-progenitor subset to each differentiated cell type according to published trajectories. This confirmed a directional loss of total enhancers during progenitor differentiation with the erythroblasts showing the greatest overall loss of enhancers (**Fig. 6C**). However, this directional loss was largely restricted to primed enhancers with active enhancers staying either at the same frequency or at a frequency that increased with differentiation (**Fig. 6C)**. The majority of primed enhancers (>90%) and surprisingly active enhancers (>72%) found in mature cells were already evident in their proximal progenitor populations (**Fig. 6D, E** and **S6D**). A significant fraction (>80%) of the active enhancers in the differentiated cell types were also already primed across the progenitor subsets (**Fig. S6E** and **F**), as noted earlier for similar subsets of mouse hematopoietic cells^9^. Interestingly, genes associated with enhancers found to be active in monocytes and erythroblasts, but already active in their corresponding progenitors produced significantly higher levels of transcripts in the mature cells compared to their inferred parental progenitors (pairwise Wilcoxon signed-rank test *p* <2×10^−5^, **Fig. 6F** and **G**). These genes were enriched in terms related to specific hematopoietic differentiation programs (Benjamini corrected *p* <10e^-30^). Strikingly, CpGs within these active enhancers showed significantly higher levels of methylation in progenitors compared to mature cells (2-sided t-test *p* < 2× 10^−16^) suggesting that CpG methylation states within active enhancers may be predictive of activity and coordinately regulated during hematopoietic differentiation. For example, *SPI1* and *EPB41* were associated with methylated active enhancers in progenitor cells that were hypomethylated in lineage-restricted cells concomitant with their expression (**Fig. 6H** and **I**). Collectively, these observations suggest that a majority of active enhancers driving terminal hematopoietic differentiation programs appear in early progenitors but constrained by CpG methylation until lineage specific programs are initiated.

To further understand how active enhancer states differ between progenitors and terminally differentiated cells, we leveraged self-organizing maps (SOM) and used ranked normalized H3K27ac density across all hematopoietic enhancers to identify progenitor, monocyte, erythroid, B and T cell enhancer clusters (methods; **Fig. S7C**). Relative to the CD34+CD38-population, GMPs and MEPs showed progression towards, and enrichment in, enhancers belonging to the monocyte and erythroblast enhancer clusters, respectively (**Fig. S7D**). Enrichment of these differentiated enhancer clusters in MEPs and GMPs supports a model where pre-existing cell type-specific active enhancer signatures emerge in proximal progenitors and may thus contribute to driving a transcriptional program activated at later stages of differentiation. Interestingly, CMPs show enrichment in both lymphoid and monocyte active enhancer clusters consistent with emerging evidence suggesting that CMPs comprise multiple subsets with different mature output capabilities^16,17^.

We next used the ROSE RANK algorithm^31^ to identify the high amplitude active enhancers and clusters of H3K27ac (super-enhancers) in each of the 8 cell types analyzed. We observed a greater than 2-fold increase in the number of super-enhancers in the mature cell types compared to the numbers of these in the progenitor populations (**Fig. 7A**). Surprisingly, we also found that a majority (>70%) of the active enhancers seen only in the mature cells were located within the boundaries of super enhancers, including the human counterparts of enhancers previously identified in differentiating mouse hematopoietic cells (**Fig. 7B** and **C**). Pathway analysis of genes associated with monocyte and erythroblast super-enhancers were enriched in terms related to leukocyte activity and erythroid differentiation, respectively (**Fig. S7E** and **F**). Taken together, these data suggest a model in which a majority of active enhancers are initially marked in progenitor populations and then further reinforced to form super-enhancers during the final processes of terminal blood cell differentiation.

**Figure 7.**
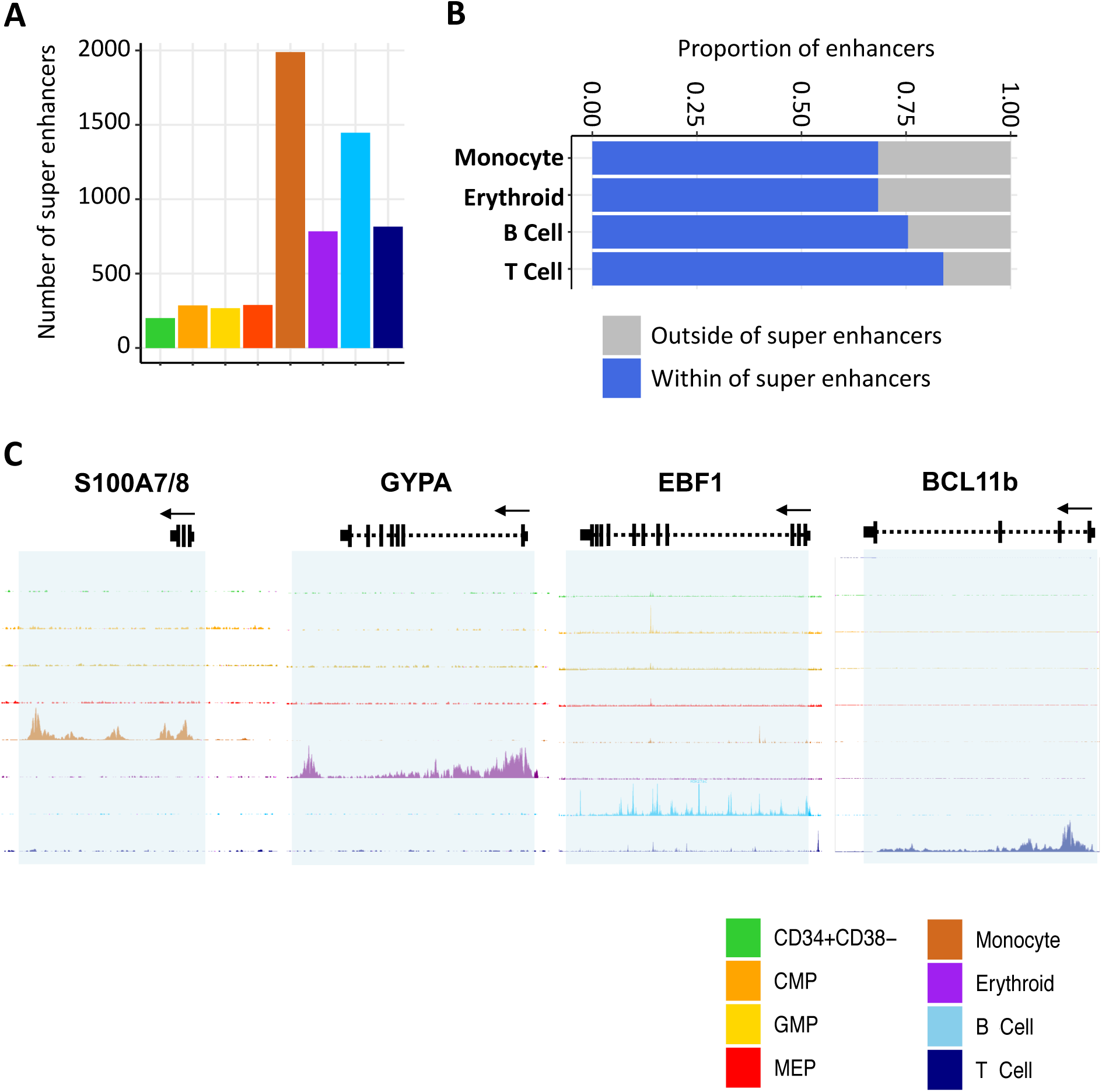
The majority of active enhancers appearing first in differentiated cell populations contribute to the establishment of super enhancers. **A)** Total number of super enhancers across cell types as indicated by the color legend. **B)** Fraction of cell type-specific active enhancers that reside within super enhancers. **C)** Genome browser view of H3K27ac density across cell types as indicated by colour at lineage-specific super enhancers indicated by the shaded box.

### EZH2 inhibition differentially alters the production of different hematopoietic lineages consistent with their acquired epigenomic features

The broad H3K27me3 domains shared by CB progenitors that are no longer present in monocytes and erythroblasts but persist in mature lymphoid cells (**Fig. 3**) led us to hypothesize that these differences have functional roles in their differentiation. To test this possibility, we first asked how the mature cell outputs of a CB cell type with dual and almost exclusive granulopoietic and B-lymphopoietic differentiation potential^16^ would be affected by exposure to either of 2 inhibitors – one, EPZ-6438, that targets EZH2, a major component of the polycomb repressive complex-2 (PRC2), and another, GSK-J4, that targets a H3K27me3 demethylases. Accordingly, CD34+CD38midCD71-CB cells were incubated with these inhibitors (0.1% DMSO as a control) for 3 weeks in cultures optimized to support their differentiation into mature GM and B cells. Analysis of the numbers of these generated in bulk cultures showed a selective decrease in CD19+ B-lineage cells in the presence of EPZ-6438 (2-sided t-test *p*<0.014), with no effects of either inhibitor on the output of cells expressing phenotypic markers of monocyte/macrophages or neutrophils (**Fig. 8A** and **B**). Additional clonal cultures showed the effect of EPZ-6438 was not explained by any toxicity to the starting cells as the total yield of clones was unaffected (118 vs 128 of 192 cells tested) with a selective reduction in the proportion with detectable CD19+ B-lineage outputs (27% vs 42% in the EPZ-6438 vs control cultures, *p*<0.001, 2-sided t-test, **Fig. 8C, S8A** and **B)**. These results indicate that continued maintenance of H3K27me3 by PRC2 is required for progenitors with lymphoid potential to differentiate into mature B-lineage cells.

**Figure 8.**
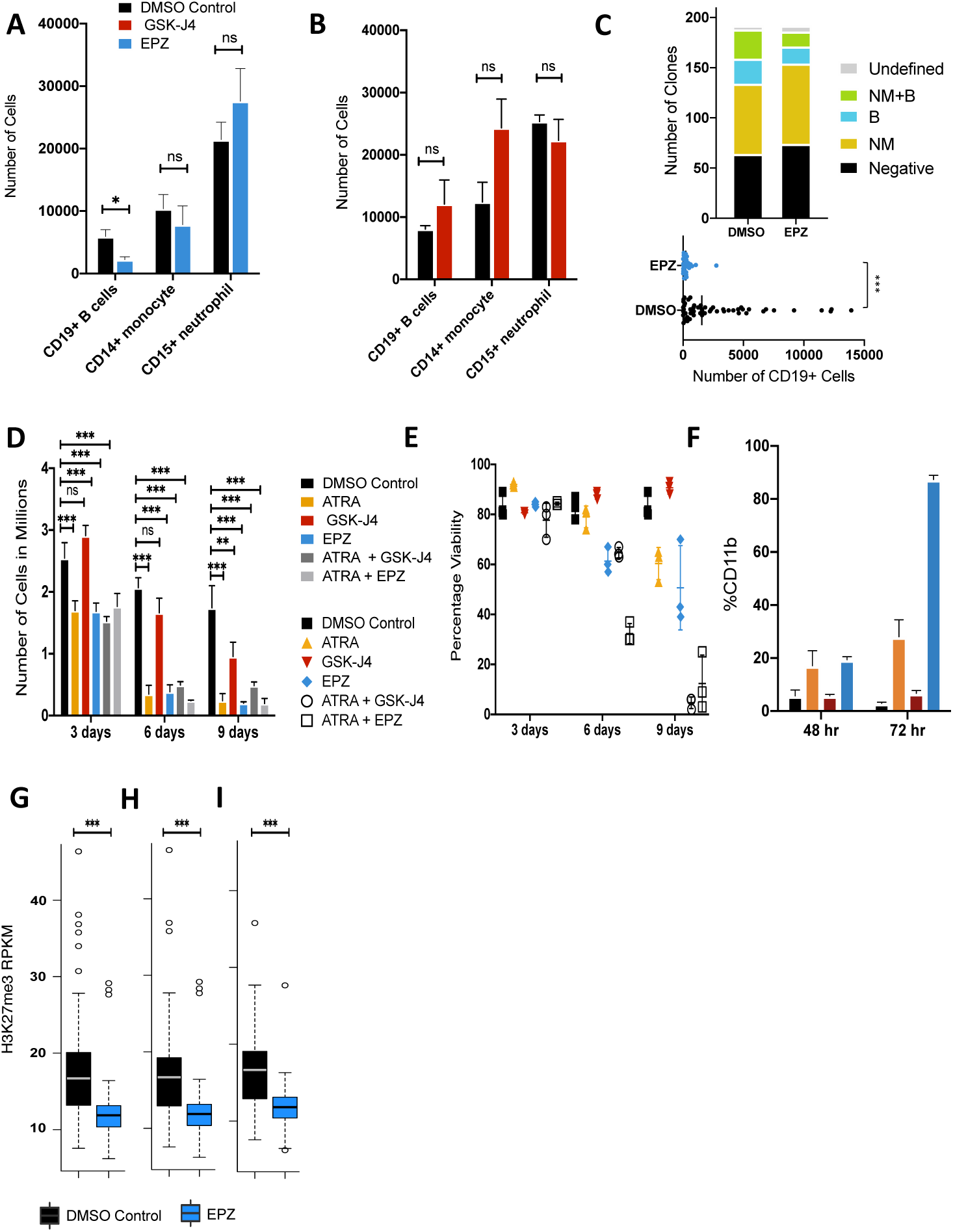
EZH2 inhibition arrests B-lymphoid cell differentiation. Total cell number of CD19+ B cells, CD14+ monocytes, and CD15+ neutrophils after 3 weeks of culturing CB-derived CD45highCD34highCD38midCD71-CD10-(P-NML) cells with lymphoid and neutrophil/monocyte lineage-stimulatory cytokines in the presence or absence of EPZ (EZH2 inhibitor) **(A)** or GSK-J4 (K27me demethylase inhibitor). **(B). C)** Bar plot showing the number of clones with different contents (top panel). M = CD14+ monocytes, N= CD15+ neutrophils, B = CD19+ B cells, negative <10 CD45+ events, undefined = no detectable mature cells. Total number of cells expressing CD19+ (bottom panel). **D)** Total number of HL60 cells in culture after 3, 6 and 9 days of treatment with ATRA, GSK-J4, or EPZ. **E)** Percentage of viable cells in **(D). F)** Percentage of CD11b+ cells assessed by FACS after 48 and 72 hours of treatment with ATRA, GSK-J4 or EPZ. H3K27me3 density at LOCKs identified in HL60 cells. **(G)** LOCKs identified in HL60 cells that overlapped with primary CB progenitor LOCKs **(H)** and were lost in monocytes **(I)**. (* *p* < 0.05, ** *p* < 0.01 and *** *p* < 0.001).

We then asked if these 2 inhibitors also affect the ability of HL-60 cells to activate a granulopoietic differentiation program^32^. After 3 days of treatment with EPZ-6438, HL60 cells showed the same growth-arrest obtained with all-trans retinoic acid (ATRA) (**Fig. 8D** and **E**) and an expected subsequent loss of viability. In contrast, in the presence of GSK-J4, the results were indistinguishable from those of the DMSO-treated controls. Confirmation of induced granulopoietic differentiation in the presence of ATRA and EPZ-6438 was obtained by FACS detection of the appearance of increased numbers of CD11b+ cells after 48 hours of exposure to these treatments compared to controls (2-sided t-test, *p* < 0.001, **Fig. 8F**). Thus, the inferred EPZ-6438-mediated removal of H3K27me3 marks from HL60 cells appears to promote the same differentiation alterations as ATRA, consistent with the loss of H3K27me3 seen to distinguish CB monocytes from their co-isolated progenitors^33^. In further support of this inference was the finding of a significantly lower H3K27me3 density (2-sided t-test, *p* < 0.001) in the EPZ-6438 treated HL60 cells compared to the DMSO control at HL60-defined LOCKs (**Fig. 8G**) in parallel with those that overlapped with LOCKs in CB progenitors (**Fig. 8H**) and no longer evident in monocytes (**Fig. 8I**).

Together these results suggest that a reduction of H3K27me3 density at LOCKs is required for the differentiation of mature myeloid cells from progenitors with that potential, whereas maintenance of H3K27me3 domains is required for the production of mature lymphoid cells.

## Discussion

Epigenetic mechanisms have long been postulated to play a central role in hematopoietic cell fate decisions^34–39^ and repressive H3K27me3 chromatin modifications have been implicated from investigations of the regulation of hematopoiesis both *in vivo* and *in vitro*^40–44^. Manipulation of H3K27me3 has been implicated particularly as impacting the steps progenitors undergo to produce mature myeloid and lymphoid cell types^30,42^ and both overexpression and inactivation of PRC2 components (responsible for the methylation of H3K27) have been reported in hematopoietic malignancies, suggesting a critical role of H3K27me3 in regulating normal hematopoiesis^45,46^. Here we revealed a broad and stable repressive H3K27me3 landscape across multiple normal progenitor subsets present in normal human CB; i.e., those conventionally defined phenotypically as an HSC-enriched subset, CMPs, GMPs and MEPs. This finding contrast markedly with the dynamic and cell-type specific landscape we report for active histone modifications. We also identified a striking and lineage-selective genome-wide H3K27me3 signature evident in 2 mature myeloid cell types (monocytes and erythroblasts) not present in 2 mature co-isolated B cells and T cells. Intriguingly, the H3K27me3 signature common to the monocytes and erythroblasts present in these CB samples also displayed a punctuated H3K27me3 profile reminiscent of that previously uniquely associated with ESCs^47^.

In the classical Waddington^48^ view, the chromatin of very primitive cells is now envisaged to exist in a more plastic state allowing for a more heterogeneous dynamic remodeling process that culminates in lineage restriction and subsequent activation of a differentiation program. One early finding in support of such privileged chromatin in ESCs is their bivalent state^21,47,49,50^ in which the nucleosomes in the promoters of developmentally important genes are marked by both permissive and repressive histone modifications (H3K4me3 and H3K27me3, respectively) that subsequently resolve to an homogenously active or repressed chromatin state as differentiation occurs^47,49^. However, subsequent epigenomic studies across a broad range of primary tissues have revealed that bivalent promoters are not unique to ESC chromatin and can be found in many cell types including fully functional, terminally differentiated cell types^3,19^. Another prevailing model of H3K27me3 occupancy during ESC differentiation posits that, upon differentiation, H3K27me3 genomic occupancy spreads outwards from discrete focal regions to occupy large genomic regions^21^, a feature that has been correlated with the differentiation capacity of the cells. Here, we demonstrate the surprising finding that H3K27me3 reverts back to a punctuated profile in cells that have differentiated but just into certain types of mature blood cells. This observation now raises the possibility that an overall loss of repressive chromatin rather than its further compaction was critical for the activation of certain (myeloid) differentiation programs. We correlated H3K27me3 contraction with a marked reduction in expression of the PRC1 complex member *BMI1*, in both monocyte and ESC cells, and demonstrated that loss of its expression and/or activity results in a genome-wide loss of H3K27me3 and certain myeloid phenotypes consistent with the model of lineage restriction in the neonatal human hematopoietic system we now propose. Why B and T cells retain a broader H3K27me3 landscape also raises interesting questions. One possibility is that this could be related to their requisite ability to activate a “stem-like” state upon antigenic stimulation in order to generate a large expansion of their progeny^51^, a feature not shared by cells within the myeloid lineages.

We also found that the observed contraction of H3K27me3 domains evident in monocytes and erythroblasts includes a specific loss of H3K27me3 marked LOCKs that in the progenitors and mature lymphoid cells show co-occupancy with another suppressive histone modification, H3K9me3. The lineage-specific restructuring of H3K27me3 specifically during differentiation into the monocyte and erythroid lineages highlights the importance of higher order chromatin structure in differentiation and establishment of cellular identity in the human hematopoietic system. H3K27me3- and H3K27me3/K9me3-enriched regions common to progenitor and lymphoid cells and lost in different mature myeloid cell types are strongly enriched in LADs, reinforcing the concept that myeloid cells can also be distinguished from lymphoid and progenitors based on lamin distribution and nucleus rigidity^52^. Taken together, these observations provide molecular support of the observation that manipulation of lamin expression specifically modulates myeloid, but not lymphoid, cell differentiation^52^.

In contrast to H3K27me3, we found the active enhancer mark H3K27ac is dynamic across all progenitor populations analyzed and the majority of active enhancers identified in the mature cells were first detected in their traditionally defined progenitor populations despite recent evidence of the considerable heterogeneity in differentiation potential and other molecular features they display^16^. Key lineage-specific regulators were identified among the genes associated with active enhancers in the progenitor populations. During differentiation, these active enhancers increase in width and amplitude and expression of associated genes increases. Our results thus suggest that priming of key regulatory regions during hematopoietic differentiation occurs in the context of traditionally described active enhancers whose activity is reinforced and increases as part of the lineage restriction process. This finding refines the currently proposed primed (H3K4me1) to an active enhancer model^9^ and suggests that additional features beyond the presence of H3K27ac may be required for full enhancer activity; for example, loss of DNA methylation at active enhancers as shown here and the formation of lineage-specific phase separated condensates^53^.

## Methods

### Preparation of human CB cells

Anonymized consented anticoagulated samples of normal CB cells were obtained with informed consent according to University of British Columbia Research Ethics Board approved protocols. CD34+ cells were enriched by EasySep™ from the light-density fraction of Lymphoprep™ or RosetteSep™-depleted CD11b+;CD3+;CD19+ cells (STEMCELL Technologies) and then used after cryopreservation in DMSO and fetal bovine serum (FBS, STEMCELL Technologies). Three pools of cord bloods consisting of collections from 646, 255 and 6 individual donors were used to isolate 10,000 cells of each compartment for histone modification ChIPseq, whole genome DNA methylation and RNAseq analyses. For the inhibitor experiments, cells were isolated from a pool of cord bloods consisting of 3 donors.

### Isolation of human CB populations

Frozen CB cells were thawed by drop-wise addition to Iscove’s Modified Dulbecco’s Medium (IMDM) (STEMCELL Technologies) supplemented with 10% FBS and 10 µg/mL DNase (Sigma-Aldrich). Cells were suspended in Hanks Balanced Salt Solution (HBSS) supplemented with 5% human serum and 1.5 µg/mL anti-human CD32 antibody (Clone IV.3; STEMCELL Technologies) and then stained with designated antibodies for 1-2 hours on ice prior to sorting on a Becton Dickinson FACSAria™ Fusion or FACSAria™ III sorter. Cells from the following populations were sorted directly into DNA LoBind tubes (Eppendorf) containing HBSS + 2% FBS: CD34+CD38-, CD34+CD38+CD10-CD7-CD135-CD45RA-(MEP), CD34+CD38+CD10-CD7-CD135+CD45RA-(CMP), CD34+CD38+CD10-CD7-CD135+CD45RA+ (GMP), CD45-CD34-GPA+ (Erythroid Precursors), CD45+CD34-CD11b+CD33+CD14+ (Monocytes), CD45+CD34-CD11b-CD33-CD19+CD7-(B cells). Naïve CD4 T cells were isolated from CB mononuclear cells (CBMCs) obtained using Lymphoprep (StemCell Technologies Inc., Canada), followed by a first step of negative selection by magnetic bead separation using EasySep Human Naïve CD4 T cell isolation kit (STEMCELL Technologies Inc., Canada) and FACS on a BD FACSAriaTM II using the following markers/staining: anti-CD3 PE (clone UCHT1; BD Bioscience), anti-CD4 BV605 (clone OKT4; BioLegend,), anti-CD25 PE-Cy7 (clone M-A251; BD Bioscience), anti-CD45RO Alexa Fluor700 (clone UCHL1; BioLegend), anti-CCR7 Alexa Fluor 647 (clone 3D12; BD Bioscience, ON, Canada), and anti-CD235 eFluor 450 (clone 6A7M eBioscience). Isolated cells were then centrifuged (500 rcf, 6 min) and had their supernatant removed prior to rapid freezing using dry ice or liquid nitrogen then stored at -80°C.

### RNA-seq

Total RNA was rRNA depleted using NEBNext rRNA Depletion Kit (New England BioLabs, E6310L). 1^st^ strand cDNA was generated using Maxima H minus First Strand cDNA Synthesis Kit (Thermo Scientific, K1652) with the addition of 1ug of Actinomycin D (Sigma, A9415). The product was purified using in-house prepared 20% PEG, 1M NaCL Sera-Mag bead solution at 1.8X ratio and eluted in 35 µL of Qiagen EB buffer. Second Strand cDNA was synthesized in a 50 µL volume using SuperScript Choice System for cDNA Synthesis (Life Technologies, 18090-019) with 12.5 mM GeneAmp dNTP Blend with dUTP. Double stranded cDNA was then purified with 20% PEG, 1M NaCL Sera-Mag bead solution at 1.8X ratio, eluted in 40 µL of Qiagen EB buffer, and fragmented using Covaris E220 (55 seconds, 20% duty factor, 200 cycles per burst). Sheared cDNA was End Repaired/Phosphorylated, single A-tailed, and Adapter Ligated using custom reagent formulations (New England BioLabs, E6000B-10) and in-house prepared Illumina forked small adapter. 20% PEG, 1M NaCl Sera-Mag bead solution was used to purify the template in-between each of the enzymatic steps. To complete the process of generating strand directionality, adapter-ligated template was digested with 5 U of AmpErase Uracil N-Glycosylase (Life Technologies, N8080096). Libraries were then PCR amplified and indexed using Phusion Hot Start II High Fidelity Polymerase (Thermo Scientific, F 549-L). An equal molar pool of each library was sequenced on HiSeq2000 (Illumina) PE75. Libraries were sequenced on a HiSeq 2500 platform following the manufacture’s protocols (Illumina, Hayward CA.). Sequence reads were aligned to a transcriptome reference generated by JAGuaR (version 2.0.2)^54^ using the GRCh37-lite reference genome supplemented by read-length-specific exon–exon junction sequences (Ensemble v75 gene annotations) and bam files were generated using Sambamba (version 0.5.5)^55^. The resulting bam files were repositioned to GRCh37-lite by JAGuaR. To quantify exon and gene expression reads per kilobase per million mapped reads (RPKM) metrics was calculated. A standardized analytical pipeline introduced by Canadian Epigenetics, Environment, and Health Research Consortium [CEEHRC] and International Human Epigenomic Consortium (IHEC) (http://ihec-epigenomes.org/) was applied to qualify the resulting data. Pairwise differential expression analysis was carried by an in-house MATLAB script, DEFine, on RPKM values that are transcripts GC biased corrected. MetaScape version 2.0^56^ (http://metascape.org) was used for genome ontology analysis of differentially expressed genes. All figures were generated by R statistical software^57^. RNA-seq data set for mouse hematopoietic cells were obtained from Lara-Astiaso, D. *et al*.^15^

### Low input native ChIP-seq

ChIP-seq was performed as previously described^18^. In brief, cells were lysed in 0.1 % Triton X-100 and Deoxycholate with protease inhibitor cocktail. Chromatin were digested by Microccocal nuclease (MNase, New England BioLabs, M0247S) at room temperature for 5 minutes and 5ul of 0.25 mM EDTA was used to stop the reaction. Antibodies against epitopes of interest, H3K4me1 (Diagenode: Catalogue# pAb-037-050, lot# A1657D), H3K27me3 (Diagenode: Catalogue# pAb-069-050, lot# A1811-001P), H3K9me3 (Diagenode: Catalogue# pAb-056-050, lot# A1675-001P), H3K4me3 (Cell Signaling Technology: Catalogue# 9751S, lot# 8), H3K27ac (PMID: 18227620), and H3K36me3 (PMID: 18227620) and digested chromatin were incubated were incubated with anti-IgA magnetic beads (Dynabeads,Themo Finsher, 10001D) for 2 hours. Pre-cleared chromatin was incubated with antibody-bead complex overnight in IP buffer (20 mM Tris-HCl pH 7.5, 2 mM EDTA, 150 mM NaCl, 0.1 % Triton X-100, 0.1 % Deoxycholate) at 4 °C. IPs were washed 2 times by Low Salt (20 mM Tris-HCl pH 8.0, 2 mM EDTA, 150 mM NaCl, 1 % Triton X-100, 0.1 % SDS) and High Salt (20 mM Tris-HCl pH 8.0, 2 mM EDTA, 500 mM NaCl, 1 % Triton X-100, 0.1 % SDS) wash buffers. IPs were eluted in elution buffer (1 % SDS, 100 mM Sodium Bicarbonate) for 1.5 hours at 65 °C. Histones were digested by Protease (Qiagen 19155) for 30 minutes at 50 C and DNA fragments were purified using Sera Mag magnetic beads in 30% PEG. Illumina sequencing libraries were generated by end repair, 3’ A-addition, and Illumina sequencing adaptor ligation (New England BioLabs, E6000B-10). Libraries were then indexed and PCR amplified (10 cycles) and sequenced on Illumina HiSeq 2500 sequencing platform following the manufacture’s protocols (Illumina, Hayward CA.).Sequence reads were aligned to GRCh37-lite using Burrows-Wheeler Aligner (BWA) 0.5.7.^58^ and converted to bam format by Sambamba (version 0.5.5). Sequence reads with BWA mapping quality scores <5 are discarded and reads that aligned to the same genomic coordinate were counted only once in the profile generation. A standardized analytical pipeline developed by the International Human Epigenomic Consortium (IHEC) (http://ihec-epigenomes.org/) was applied to qualify the resulting data.

### ChIP-seq analysis

Genome browser tracks were generated by converting bam files to wiggle files using a custom script (http://www.epigenomes.ca/tools-and-software). Wiggle files then converted to bigwig for display on UCSC genome browser by UCSC tool, wig2Bigwig script. MACS2^59^ was employed to identify enriched regions with a false discovery rate (FDR) value of ≤ 0.01 for H3K4me3, H3K4me1 and H3K27ac peaks in ChIP-seq data. Finder2.0 (http://www.epigenomes.ca/tools-and-software/finder) was used to identify enriched regions for H3K27me3 and H3K9me3 ChIP-seq data. Enriched regions overlapping with ENCODE blacklist regions were eliminated. ChIP-seq signal was calculated as tag density generated using HOMER v4.10^60^ and normalized to total number of tags within enriched regions. Heat maps of fragment distribution at promoters were generated by deeptools^61^. All other figures were generated by R statistical software^57^. ChromHMM^62^ was used to identify 18 chromatin states based on MACS2^59^ identified enriched regions in each cell type as previously described^23^. To identify active enhancers, we first found MACS2 identified enriched H3K27ac regions in all populations (CD34+ CD38-, CMP, GMP, MEP, Monocyte, Erythroid, B cell and T cell) and created an enhancer catalogue of hematopoietic cells. Any enriched region that overlap with +/-2Kb of TSS defined by Ensemble v75 human gene annotation was discarded. All H3K27ac libraries were subsampled to the depth of the library with the lowest depth. Next, enhancers with H3K27ac tags > 40 in at least one cell type were selected. Then normalized signal H3K27ac tag density was calculated across all regions in our enhancer catalogue for each cell type and enhancers with signal >25 were marked as active in the corresponding cell type. Same strategy was used to identify poised enhancers with H3K4me1 enriched regions with the exception of signal threshold of 40. SOM analysis was carried by oposSOM^63^ R package on ranked normalized signal of H3K27ac at union of H3K27ac enriched regions identified across all cell types. Enhancer gene association was done by bedtools^64^ closestBed command with default parameters using Ensemble v75 human gene annotation. Promoters (+/2kb of TSS of protein coding) with distance of greater than 50Kb to enhancer regions were eliminated. We utilize GREAT, an online ontology analysis tool, for gene ontology analysis of regulatory regions. To identify H3K27me3 LOCKs, MACS2 broad mode was used to call peaks for H3K27me3 with FDR cut off of 0.05. The resulting peaks were then entered into CREAM^65^ with WScutoff of 1.5, MinLength of 1000 and peakNumMin of 2 to identify H3K27me3 LOCKs. H3K27me3 marked genes in CD34+CD38-, monocyte, erythroid precursor, B and T cells were annotated by overlapping FindER identified H3K27me3 enriched regions with hg19v75 ensemble gene annotations. We then selected for genes with >20% of their genome covered with H3K27me3.

### Whole genome bisulfite sequencing

Whole genome bisulfite libraries were constructed as previously described^66^. In brief, genomic DNA was sheered and subjected to bisulfite conversion using the MethylEdge Bisulfite Conversion kit (Promega, N1301) using a bead-based automated protocol. Bisulfite-converted DNA was mixed with 180 μl of MethylEdge Binding Buffer and 1.8μl of 20 mg/ml decontaminated MagSi-DNA all round silica beads (MagnaMedics, MD02018) and left at room temperature for 15 minutes and washed twice with 220 μl of 80% ethanol for 30 seconds. 60 μl of MethylEdge desulfonation buffer was added to the beads and incubated at room temperature for 15 minutes, then washed twice with 100 μl of 80% ethanol and dried for 1 minute. To elute the DNA, 20 μl 10 mM Tris-HCL, pH 8.5 (Qiagen, 19086) were added to DNA-bead mix and incubated in a Thermomixer C (Eppendorf, 5382000015) at 56°C while being centrifuged at 2,000 rpm for 15 minutes. The Bisulfite converted single stranded DNA was converted to double stranded DNA through 1 cycle of PCR with random hexamer as previously described^66^ followed by standard illumine library construction. Libraries were aligned using Novoalign V3.02.10 (www.novocraft.com) to human genome assembly GRCh37 (hg19). Duplicate reads were marked by Picard V1.31 (http://picard.sourceforge.net). and discarded (http://picard.sourceforge.net). Methylation calls were generated using Novomethyl V1.01 (www.novocraft.com). All figures were generated by R statistical software^57^.

### HL60 treatment with ATRA, EPZ-6438 and GSK-J4

HL60 cells were cultured at 500,000 cells/mL in RPMI (StemCell, 36750) with 10% FBS (Sigma, F1051) with 1X Penicillin-Streptomycin (Gibco, Life Technologies, Fisher Thermo. 15140122) and 1X GlutaMAX (Gibco, Life Technologies, Fisher Thermo. 35050061) in the presence of 1 µM ATRA (Sigma, R2625), 20 µM EPZ-6438 (Selleckchem, S7128) and 1 µM GSK-J4 (Sigma, 420205) in 96-well U-shape bottom plates for 9 days. Cells were treated with each compound every 3 days and reset to original seeding density. Concentration and viability were assessed every 3 days using a Countess II automated cell counter (Thermo Finsher, AMQAX100).

### Quantitative measurement of CD11b in HL60

HL60 cells were cultured at 1,000,000 cells/mL in RPMI (StemCell, 36750) with 10% FBS (Sigma, F1051) with 1X Penicillin-Streptomycin (Gibco, Life Technologies, Fisher Thermo. 15140122) and 1X GlutaMAX (Gibco, Life Technologies, Fisher Thermo. 35050061) in presence of 0.1uM ATRA (Sigma, R2625), 20uM Tazemetostat (Selleckchem, S7128) and 1uM GSK-J4 (Sigma, 420205) in 96 well V-shape bottom plate for 48 and 72 hours. Concentration of DMSO control was adjusted to maximum amount of DMSO (1.1%) used in treated wells. After 48 and 72 hours, all cells were harvested, suspended in Hanks’ Balanced Salt Solution (HBSS; STEMCELL Technologies) supplemented with 2% FBS and 1.5ug/mL anti-human CD32 antibody (Clone IV.3; STEMCELL Technologies). Cells were stained for 20-30 minutes on ice with 1:200 anti-CD11b-BV711 (Clone M1/70; Biolegend) prior to analysis by flow cytometry. Flow cytometry data were analyzed in R using the package flowCore^67^ and custom scripts.

### Stromal cell-containing cultures of CB cells

CB cells pooled from 3 donors were thawed, stained with antibodies. For “bulk cultures, 50 cells with a FACS-purified CD45highCD34highCD38midCD71-CD10-(P-NML) phenotype were deposited into each well preloaded with 100,000 MS-5 stromal cells and alpha-MEM medium with 2 mM glutamine, 7.5% FBS and 10^−4^M β-mercaptoethanol (Sigma) plus 50 ng/ml SCF (Novartis), 10 ng/ml FLT3L (Immunex), 1 ng/ml IL-3 (Novartis), and 3 units/mL EPO (STEMCELL Technologies) in the presence of 1 µM GSK-J4 (Sigma, 420205) or 0.4 µM EPZ-6438 (Selleckchem, S7128) as indicated. The same dose of inhibitor was added to each half medium change performed after 4 and 7 days, respectively. Cells were cultured with 50 ng/ml SCF, 10 ng/mL FLT3L, 10 ng/ml IL-7 (R&D Systems), 3 units/ml EPO for the second week and 10 ng/ml IL-7 and 3 units/ml EPO only for the third week. After 3 weeks, all cells were harvested, stained with antibodies, and assessed by FACS to detect cultures of >10 cells belonging to one or more of the following cell populations: monocytes (CD45+CD33+CD14+ cells), neutrophils (CD45+CD33+CD15+ cells) and B cells (CD45+CD33-CD14-CD15-CD19+ cells).

For clonal cultures, single P-NML cells were deposited into each well of a 96-well plate preloaded with 9,000 MS-5 cells and 333 each of M210B4 mouse fibroblasts expressing human IL-3 and G-CSF, and sl/sl mouse fibroblasts expressing human SCF and IL-3, and human FLT3L, with alpha-MEM medium with 2 mM glutamine, 7.5% FBS and 10−4 M β-mercaptoethanol (Sigma) plus 50 ng/mL, SCF (Novartis), 10 ng/mL FLT3L (Immunex) added for the first 2 weeks. Weekly half-medium changes were performed. After 3 weeks, all cells were harvested, stained with antibodies and assessed by FACS to detect clones of >10 monocytes, neutrophils or B cells using the same FACS analysis protocol as for the bulk cultures. Clones containing >10 CD45+ events in the absence of any mature cells were classified as “undefined”. Clones with <10 CD45+ events were classified as “negative”.

### CRISPR-KO of *BMI1*

We generated *BMI1*-KO HL60 cells using Alt-Râ CRISPR-Cas9 genome editing System following manufacturer’s guide (Integrated DNA Technologies, Inc.). Ribonucleoprotein complexes (crRNA:tracrRNA duplex) were delivered using Neonâ Transfection System. For HL60, 2’10^5^ cells per 10ul reaction were electroporated at 1600V, 10ms, 3 pulses. For DU145, 0.8’10^5^ cells per 10ul reaction were electroporated at 1250V, 20ms, 2 pulses. Samples of amplicons were generated by PCR using designed primers targeting at *BMI1* cut sites. IDT online tool was used to design the following guide RNAs (**Table S1**). Efficiency of on target CRISPR events were assessed by Alt-R genome editing detection kit (IDT). BMI1 level was assessed by western blot using standard SDS-PAGE in 4%-12% gel of NuPAGE electrophoresis system (Life Technology, Thermo Fisher). Antibodies are: β-actin (Cell Signaling: Catalogue# 13E5) and BMI1 (Cell Signaling: Catalogue# D20B7).

### Western blot

Total histone protein was extracted using Histone Extraction kit (Abcam, Catalogue#:ab113476) according to the manufacture’s protocol. Total histone protein was quantified using BCA assay. H3, H3K9me3, H3K27me3 and H2AK119Ub modified histones were analyzed by standard SDS-PAGE in 4%-12% gel of NuPAGE electrophoresis system (Life Technology, Thermo Fisher). Antibodies are: Rabbit anti-H3(N-terminal) (Sigma, Catalogue#: H9289, 1:1000), H3K9me3 (Diagenode: Catalogue# pAb-056-050 1:1000) and H2AK119Ub (Cell Signaling: Catalogue# D27C4). Mouse anti-H3K27me3 antibody, from Dr. Hiroshi Kimura (Kimura et al., 2008). Secondary antibodies are goat anti-rabbit IRDye-800CW (LI-COR, 1:10000), goat anti-mouse IRDye-680RD (LI-COR, 1:10000). The blots were imaged by LI-COR Odysseyâ CLx Infrared Imaging System. ImageJ was used for analysis of western blot images.

## Acknowledgements

The authors thank the production and technical staff at Canada’s Michael Smith Genome Sciences Centre for generating of human CB T cell reference epigenomes and their technical expertise in sequence generation and analysis. The authors also thank Darcy Wilkinson and Margaret Hale in the Eaves lab for collecting and processing CB cells, respectively. A.L., and C.H. were supported by Frederick Banting and Charles Best Canada Graduate Scholarship Doctoral Awards (CGS-D), and F.W. by a University of BC Graduate Fellowship. This work was supported by the Canadian Institutes of Health Research (EP1-120589 & CEE-151619) and Genome Canada (C41EMT &C32EMT) under the Canadian Epigenetics, Environment and Health Research Consortium (to M.H.) and by the Terry Fox Research Institute (#1039 to MH, #1074 & TFF-122869 to M.H. and C.J.E.)

## Contributions

A.L. and M.H. conceptualized the study design. A.L. performed the analysis. M.H., C.J.E., C.H., M.B., A.C., A.H.M. and J.Y.A.Z. contributed to the data analysis and interpretation. A.L., C.H., D.J.H.F.K., F.W., J.CH.W., C.Q, M.W., M.M. P.L. and Z.S. performed experimental work. A.L., M.H. and C.J.E. wrote the manuscript.

## Accession Numbers

The sequencing data reported in this study is accessible through the European Genome-Phenome Archive (EGA) under accession number EGAD00001004950.

## Notes

### Competing Interest Statement

The authors have declared no competing interest.

