## Supplemental Figures for "Polycomb contraction differentially regulates terminal human hematopoietic differentiation programs"

### Supplemental Figures and Legends

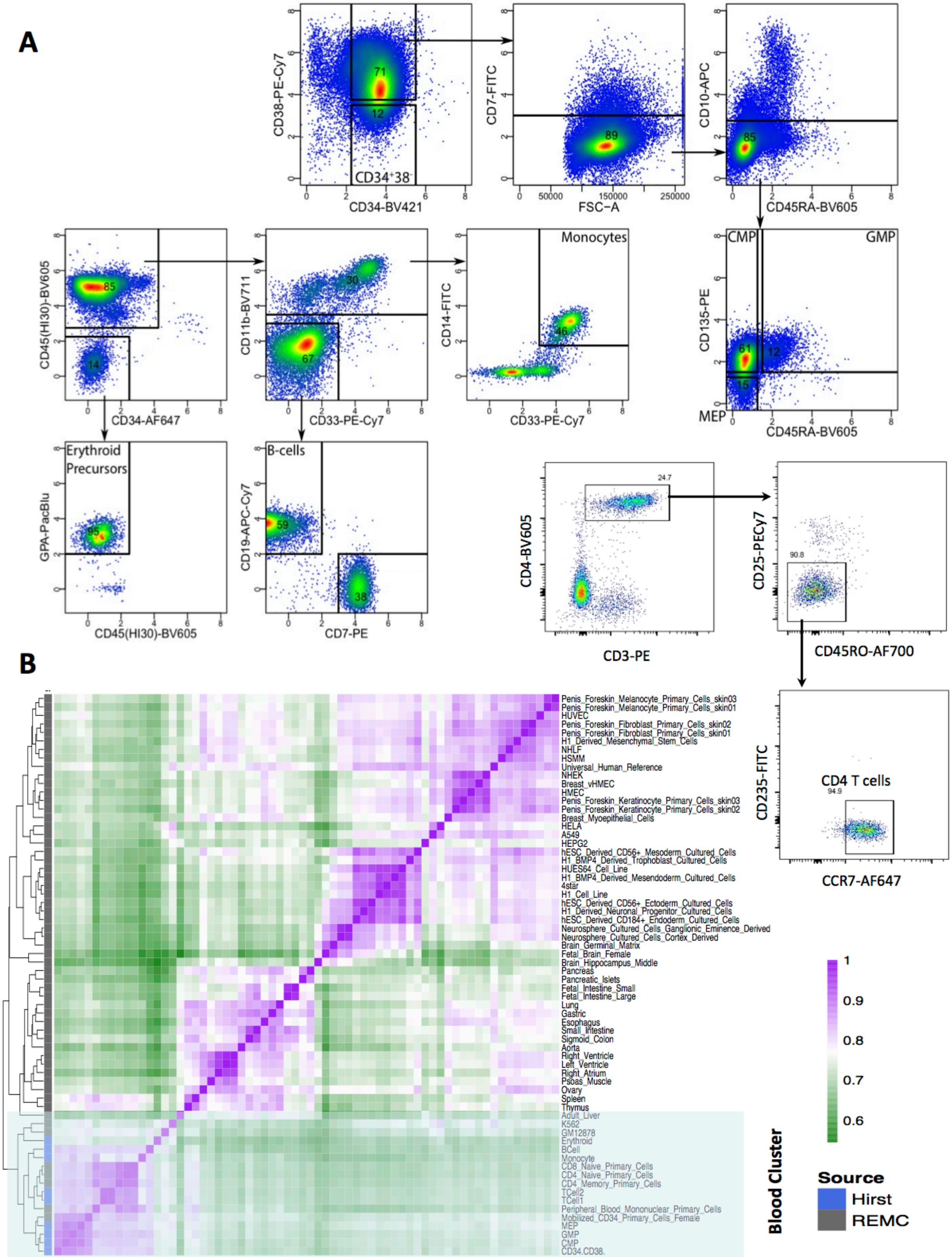

**Figure S1. Sorting strategy for hematopoietic populations profiled in this study. A)** Representative examples of sorting strategies for CD34+CD38-, CMP, GMP, MEP, monocytes, erythroblasts and B and T cell isolated from cord blood pools. **B)** Unsupervised hierarchical clustering and heatmap of pairwise spearman correlations for protein coding gene RPKM values across blood cell types profiled in this study in the context of all cell types profiled by NIH Epigenome RoadMap Consortium. The cluster of blood cell types is indicated by the shaded box.

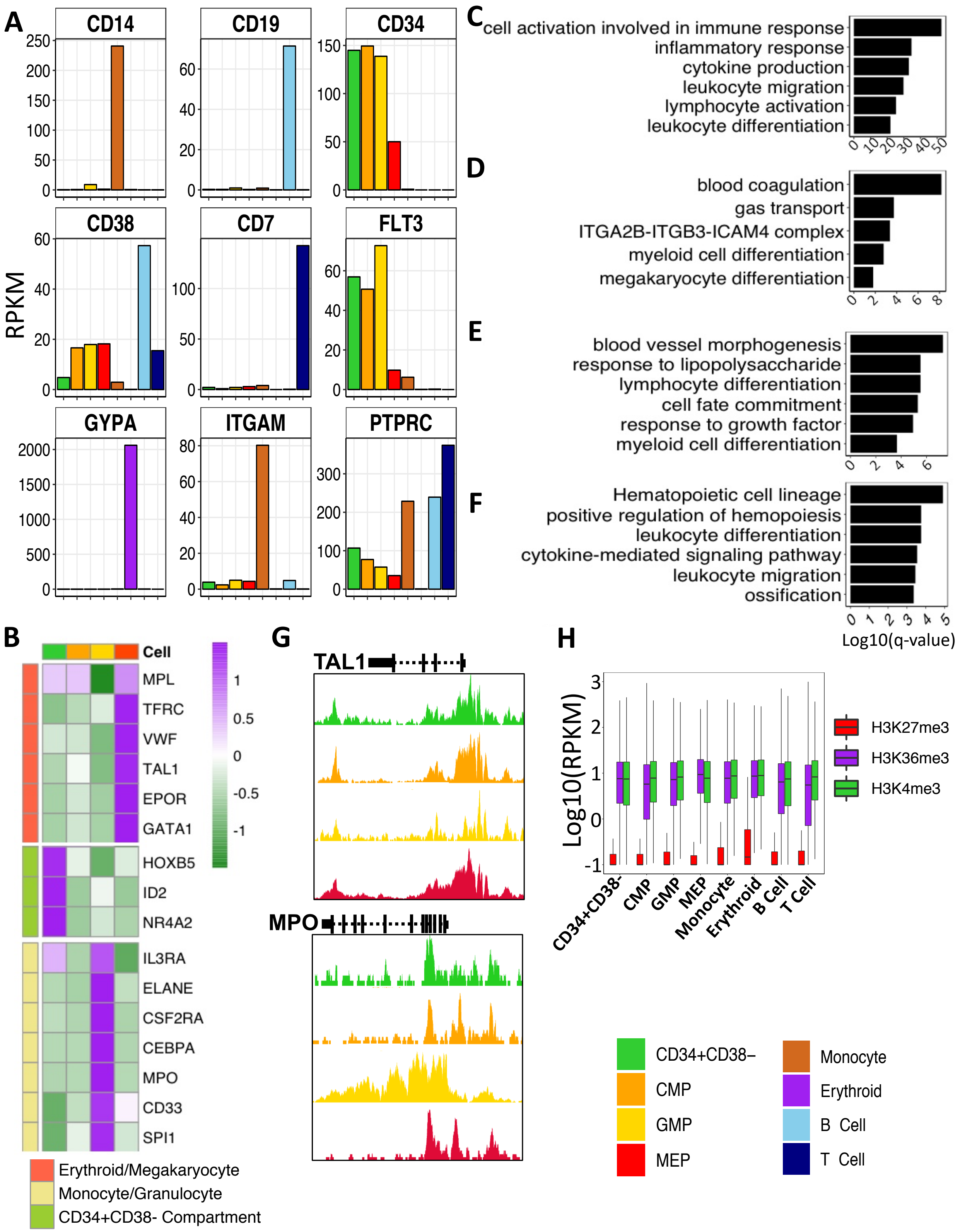
A
RPKM
CD14
CD19
CD34
CD38
CD7
FLT3
GYPA
ITGAM
PTPRC
C
cell activation involved in immune response
inflammatory response
cytokine production
leukocyte migration
lymphocyte activation
leukocyte differentiation
D
blood coagulation
gas transport
ITGA2B-ITGB3-ICAM4 complex
myeloid cell differentiation
megakaryocyte differentiation
E
blood vessel morphogenesis
response to lipopolysaccharide
lymphocyte differentiation
cell fate commitment
response to growth factor
myeloid cell differentiation
F
Hematopoietic cell lineage
positive regulation of hemopoiesis
leukocyte differentiation
cytokine-mediated signaling pathway
leukocyte migration
ossification
Log10(q-value)
B
Cell
MPL
TFRC
VWF
TAL1
EPOR
GATA1
HOXB5
ID2
NR4A2
IL3RA
ELANE
CSF2RA
CEBPA
MPO
CD33
SPI1
Erythroid/Megakaryocyte
Monocyte/Granulocyte
CD34+CD38- Compartment
G
TAL1
MPO
H
Log10(RPKM)
H3K27me3
H3K36me3
H3K4me3
CD34+CD38-
CMP
GMP
MEP
Monocyte
Erythroid
B Cell
T Cell
CD34+CD38–
CMP
GMP
MEP
Monocyte
Erythroid
B Cell
T Cell

**Figure S2. Progenitor populations possess unique expression profiles. A)** Expression (RPKM) of cell type specific cell surface marker genes across cell types as indicated by the colour legend on the bottom right. **B)** Expression of previously identified progenitor population specific genes across CD34+CD38-, CMP, GMP and MEP. Gene ontology analysis of GMP **(C)**, MEP **(D)**, CD34+CD38- **(E)**, and CMP **(F)** differentially expressed gene sets identified by DEFine (FDR > 0.01). **G)** Genome browser view of H3K4me3 density in progenitor populations at the *TAL1* and *MPO* locus across CD34+CD38-, CMP, GMP and MEP. **H)** Expression of genes marked with H3K4me3, H3K27me3 or H3K36me3 across each cell type.

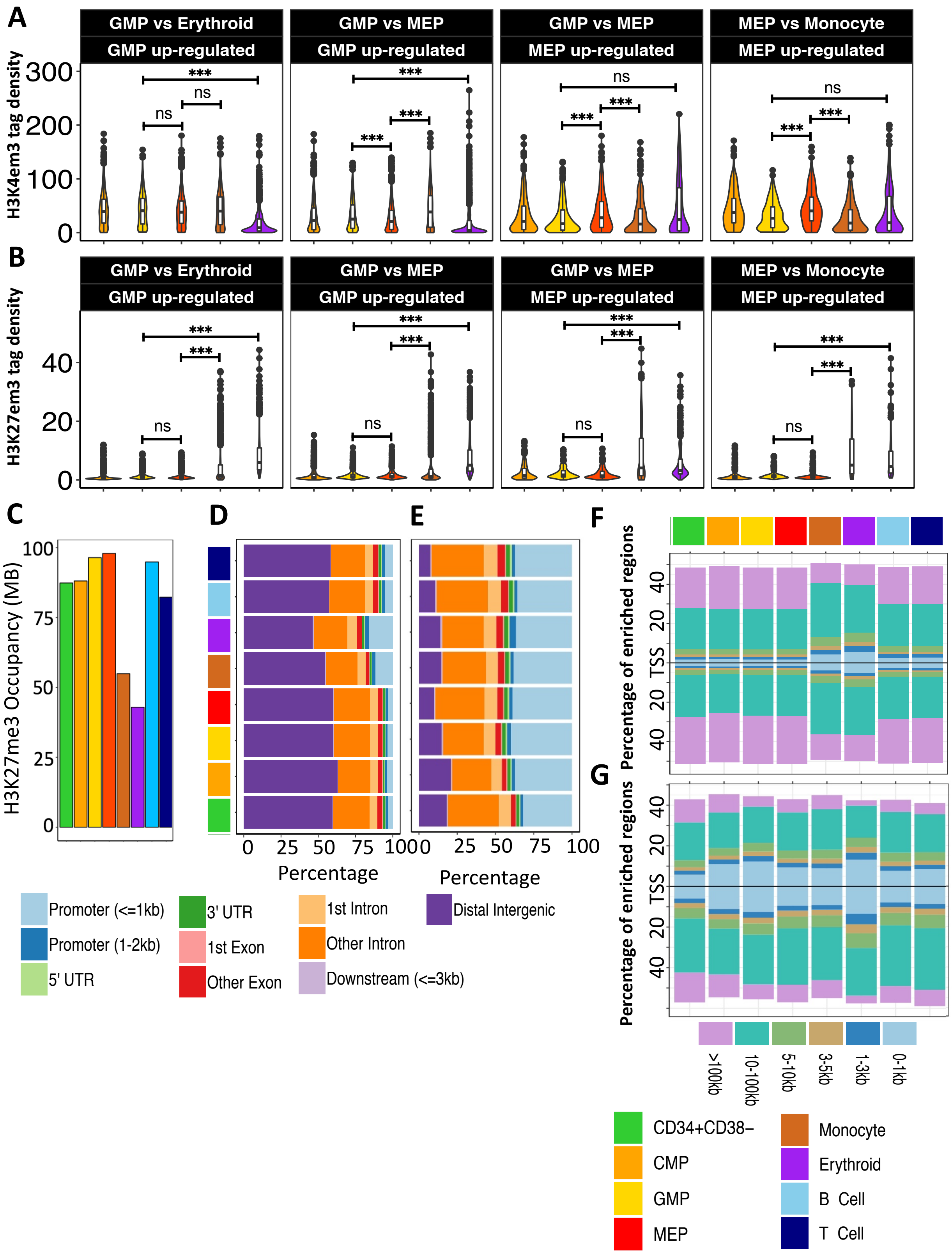
A
GMP vs Erythroid
GMP up-regulated
GMP vs MEP
GMP up-regulated
GMP vs MEP
MEP up-regulated
MEP vs Monocyte
MEP up-regulated
H3K4em3 tag density
B
H3K27em3 tag density
C
H3K27me3 Occupancy (MB)
D
E
Percentage
F
G
Percentage of enriched regions
TSS
>100kb
10-100kb
5-10kb
3-5kb
1-3kb
0-1kb
Promoter (<=1kb)
Promoter (1-2kb)
5' UTR
3' UTR
1st Exon
Other Exon
1st Intron
Other Intron
Downstream (<=3kb)
Distal Intergenic
CD34+CD38−
CMP
GMP
MEP
Monocyte
Erythroid
B Cell
T Cell

**Figure S3. H3K27me3 promoter density signatures are conserved across progenitor populations.** H3K4me3 **(A)** and H3K27me3 **(B)** tag density ±2 Kb of transcription start sites of genes up-regulated in GMP and MEP across CMP, GMP, MEP, monocytes and erythroblasts as indicated by the colour legend on the bottom right. **C)** Plot of the cumulative number of base pairs marked by H3K27me3 across cell types indicated by colour legend on the bottom right. Percentage of H3K27me3- **(D)** and H3K4me3-**(E)** enriched regions at the genomic features indicated. Percentage of H3K27me3 **(F)** and H3K4me3 **(G)** enriched regions with respect to their binned distance to the TSS of coding genes across cell types indicated by colours. Distance from TSS for each bin indicated in the bottom panel. (*** $p < 0.001$).

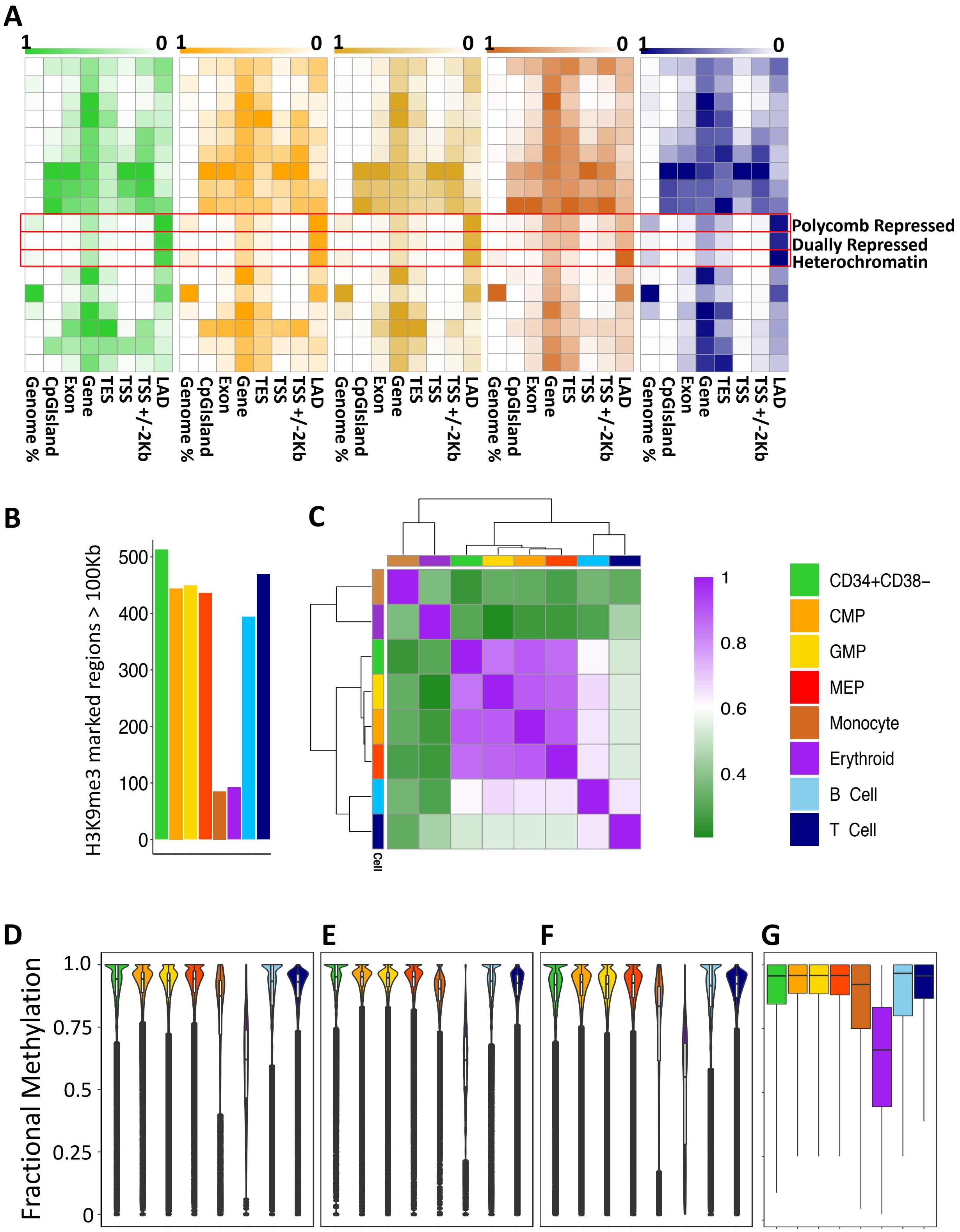

A
1
0
Polycomb Repressed
Dually Repressed
Heterochromatin
Genome %
CpGIsland
Exon
Gene
TES
TSS
TSS +/-2Kb
LAD
B
H3K9me3 marked regions > 100Kb
500
400
300
200
100
0
C
Cell
1
0.8
0.6
0.4
CD34+CD38−
CMP
GMP
MEP
Monocyte
Erythroid
B Cell
T Cell
D
E
F
G
Fractional Methylation
1.0
0.75
0.5
0.25
0

**Figure S4. ChromHMM identified polycomb repressed regions in progenitor cells are enriched in LADs**. **A**) Enrichment of ChromHMM chromatin states within genomic features across cell types as indicated by the colour legend. **B)** Plot of the cumulative number of base pairs marked by H3K9me3 across cell types indicated by colour legend **C)** Unsupervised hierarchical clustering and heatmap showing pairwise spearman correlation of H3K9me3 density genome-wide across cell types as indicated by the colour legend. A H3K27me3 LOCK (FDR < 0.05) present in progenitor cells but lost in monocytes and erythroblasts is indicated by the shaded box. Fractional CpG methylation signal within ChromHMM defined H3K9me3 **(D),** H3K27me3 **(E)** and H3K9me3/H3K27me3 (**F**) enriched regions and genome-wide (**G**) across cell types as indicated by colour legend.

.

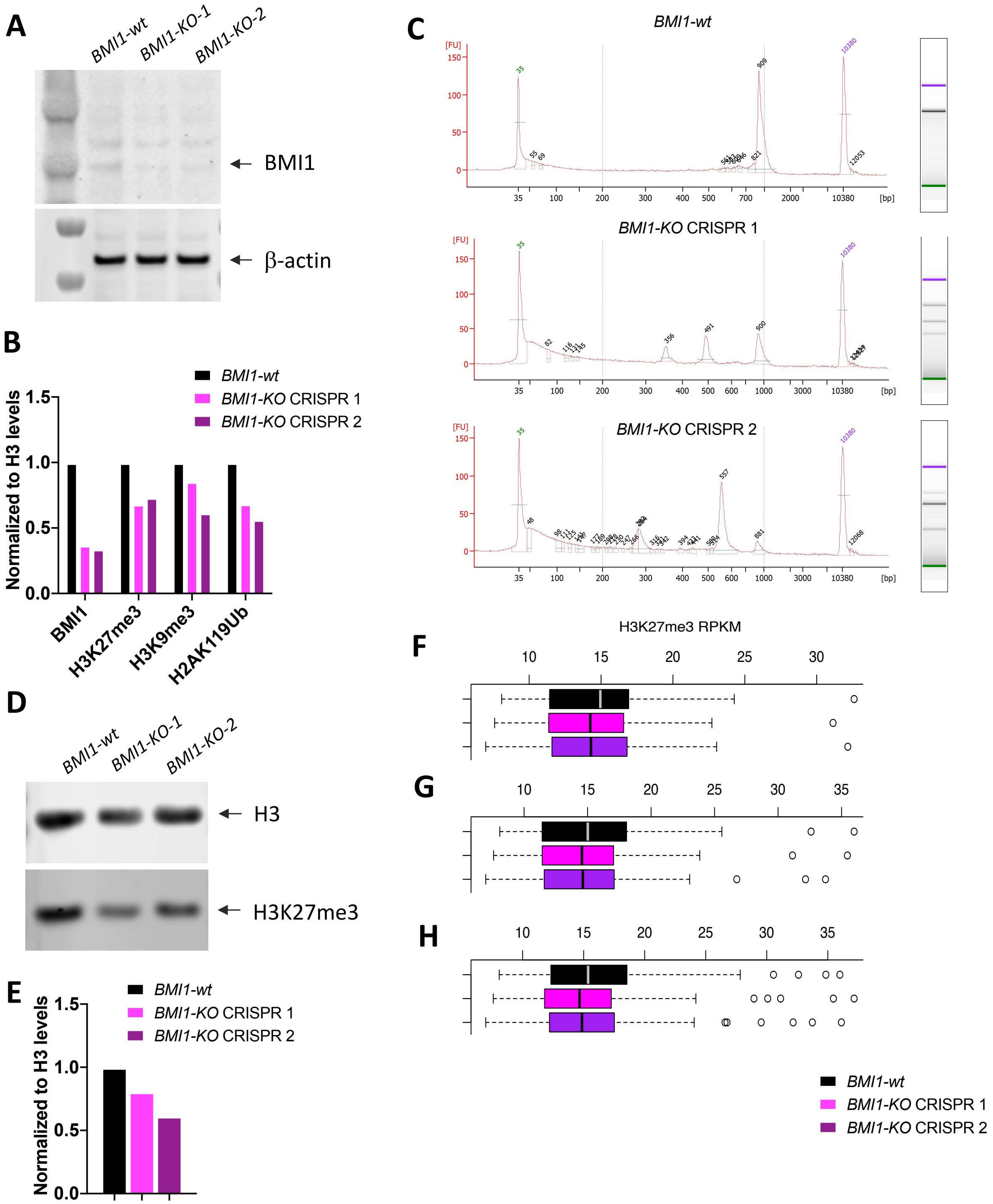
A
BMI1-wt
BMI1-KO-1
BMI1-KO-2
BMI1
β-actin
B
Normalized to H3 levels
BMI1-wt
BMI1-KO CRISPR 1
BMI1-KO CRISPR 2
BMI1
H3K27me3
H3K9me3
H2AK119Ub
C
BMI1-wt
BMI1-KO CRISPR 1
BMI1-KO CRISPR 2
[FU]
[bp]
D
BMI1-wt
BMI1-KO-1
BMI1-KO-2
H3
H3K27me3
E
Normalized to H3 levels
BMI1-wt
BMI1-KO CRISPR 1
BMI1-KO CRISPR 2
H3K27me3 RPKM
F
G
H
BMI1-wt
BMI1-KO CRISPR 1
BMI1-KO CRISPR 2

**Figure S5. BMI1 loss reduces genome-wide levels of H3K27me3. A)** Global measurement of BMI1 and β-actin by western blot across *BMI1-wt* and *KO*. **B)** Quantitative measurement of western blot intensity shown in **Figure 5C** and **(A)**. **C)** Agilent bioanalyzer profile of PCR amplicon after T7E1 mutation cleavage assay in CRISPR Control (*BMI1-wt*) and *BMI1-KO* replicates. **D)** Global measurement of H3 and H3K27me3 by western blot *BMI1-wt* and *KO*. **E)** Quantitative measurement of western blot intensity shown in **(D)**. H3K27me3 density at LOCKs identified in HL60 **(F)**, those overlapped with primary CB progenitor LOCKs **(G)** and those that are lost in monocyte **(H)**.

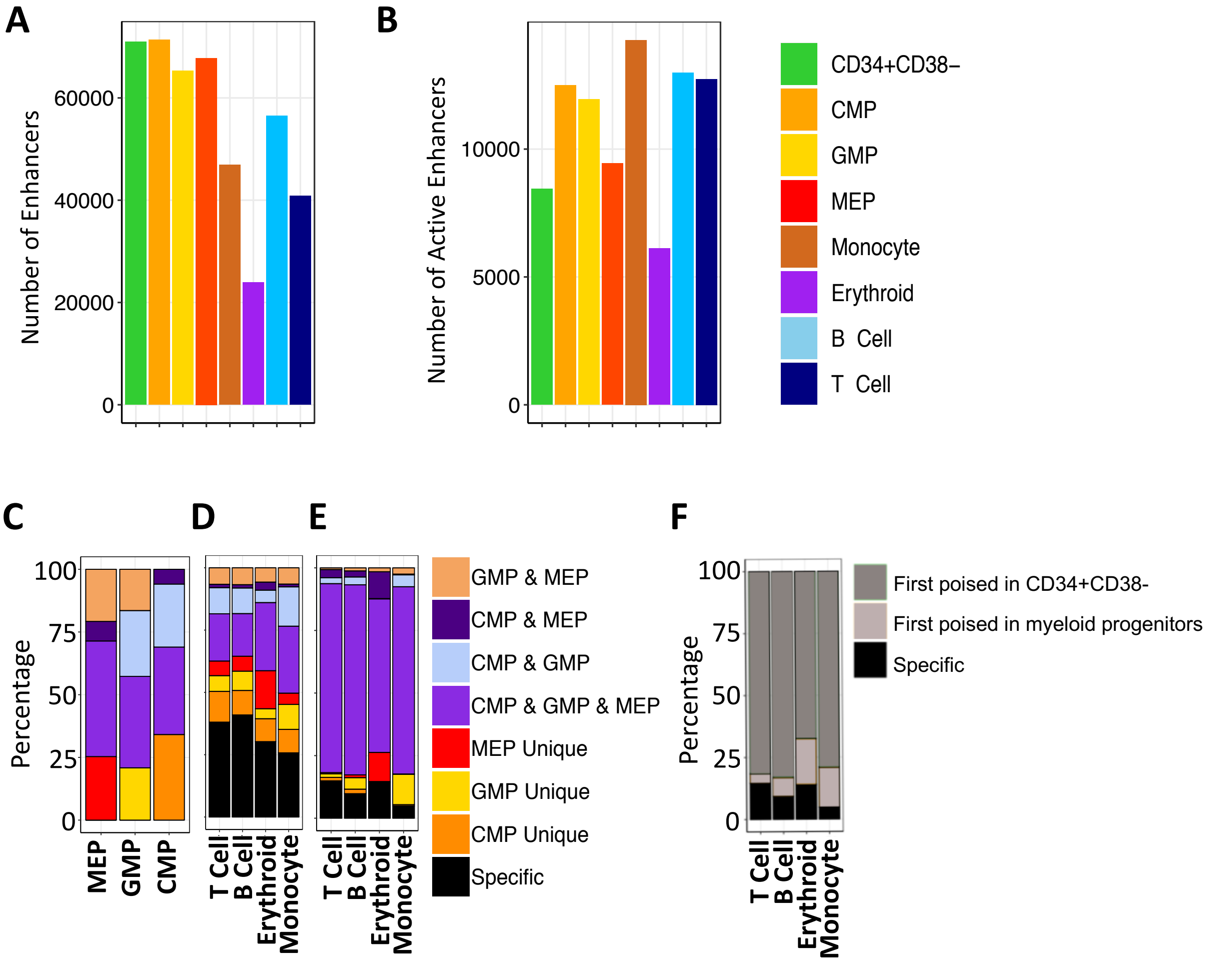
A
B
C
D
E
F
Number of Enhancers
Number of Active Enhancers
60000
40000
20000
0
10000
5000
CD34+CD38−
CMP
GMP
MEP
Monocyte
Erythroid
B Cell
T Cell
Percentage
100
75
50
25
MEP
GMP
CMP
T Cell
B Cell
Erythroid
Monocyte
GMP & MEP
CMP & MEP
CMP & GMP
CMP & GMP & MEP
MEP Unique
GMP Unique
CMP Unique
Specific
First poised in CD34+CD38-
First poised in myeloid progenitors

**Figure S6. Enhancers dynamics in erythroid precursor, monocyte and differentiated lymphoid cells. A)** Number of enhancers (active and primed enhancers) across cell types. **B)** Total number of active enhancers across cell types as indicated by the colour legend. **C)** Percentage of active enhancers that are shared between or unique to each progenitor populations. **D)** Percentage of active enhancers that are present in progenitor populations or *de novo* (specific) in differentiated cells as indicated by the colour legend. **E)** Percentage of active enhancers that are poised in progenitor populations or *de novo* in differentiated cells as indicated by the colour legends. **F)** Percentage of active enhancers present in differentiated cells that are first poised in CD34+CD38- cells, progenitor cells, or appear *de novo* in erythroblasts, monocytes or lymphoid cells. Cell types are indicated by the colour legend.

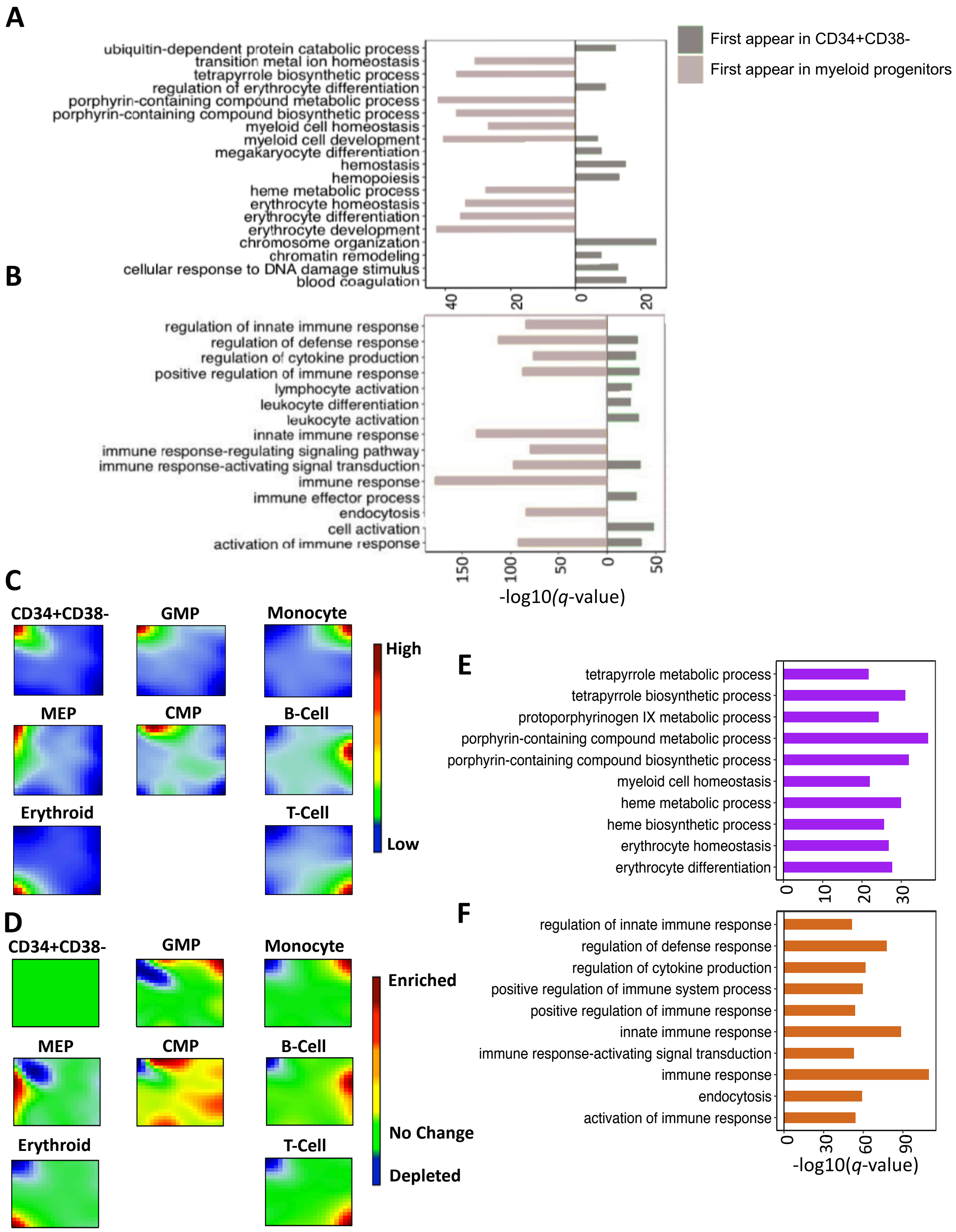
A
First appear in CD34+CD38-
First appear in myeloid progenitors
ubiquitin-dependent protein catabolic process
transition metal ion homeostasis
tetrapyrrole biosynthetic process
regulation of erythrocyte differentiation
porphyrin-containing compound metabolic process
porphyrin-containing compound biosynthetic process
myeloid cell homeostasis
myeloid cell development
megakaryocyte differentiation
hemostasis
hemopoiesis
heme metabolic process
erythrocyte homeostasis
erythrocyte differentiation
erythrocyte development
chromosome organization
chromatin remodeling
cellular response to DNA damage stimulus
blood coagulation
40
20
0
20
B
regulation of innate immune response
regulation of defense response
regulation of cytokine production
positive regulation of immune response
lymphocyte activation
leukocyte differentiation
leukocyte activation
innate immune response
immune response-regulating signaling pathway
immune response-activating signal transduction
immune response
immune effector process
endocytosis
cell activation
activation of immune response
150
100
50
0
50
-log10(q-value)
C
CD34+CD38-
GMP
Monocyte
MEP
CMP
B-Cell
Erythroid
T-Cell
High
Low
D
CD34+CD38-
GMP
Monocyte
MEP
CMP
B-Cell
Erythroid
T-Cell
Enriched
No Change
Depleted
E
tetrapyrrole metabolic process
tetrapyrrole biosynthetic process
protoporphyrinogen IX metabolic process
porphyrin-containing compound metabolic process
porphyrin-containing compound biosynthetic process
myeloid cell homeostasis
heme metabolic process
heme biosynthetic process
erythrocyte homeostasis
erythrocyte differentiation
0
10
20
30
F
regulation of innate immune response
regulation of defense response
regulation of cytokine production
positive regulation of immune system process
positive regulation of immune response
innate immune response
immune response-activating signal transduction
immune response
endocytosis
activation of immune response
0
30
60
90
-log10(q-value)

**Figure S7. Lineage-specific enhancers are marked by H3K27ac in hematopoietic progenitor subsets.** GREAT pathway enrichment analysis of active enhancers in erythroblasts **(A)** and monocytes **(B)** that are first apparent in CD34+CD38- or other progenitor subsets. **C)** SOM plot of rank normalized H3K27ac signal contained within the hematopoietic enhancer catalogue across cell types profiled in this study. **D)** SOM plot of rank normalized H3K27ac signal within hematopoietic enhancers with respect to CD34+CD38- for each cell type. Gene ontology analysis of super enhancers in erythroblasts **(E**) and monocytes **(F)**.

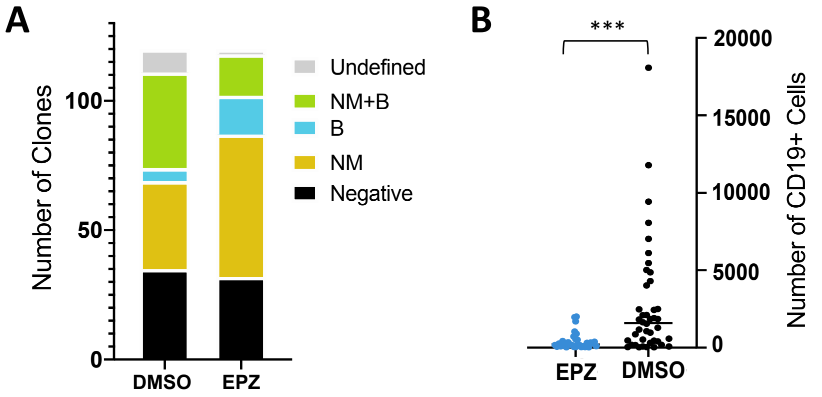
A
Number of Clones
0
50
100
DMSO
EPZ
Undefined
NM+B
B
NM
Negative
B
***
EPZ
DMSO
Number of CD19+ Cells
0
5000
10000
15000
20000

**Figure S8. EZH2 inhibition impedes B cell production. A)** Bar plot showing the number of clones with different contents (left panel). M = CD14+ monocytes, N= CD15+ neutrophils, B = CD19+ B cells, negative = <10 CD45+ events, undefined = no detectable mature cells. **B)** Total number of cells expressing CD19+ (right panel). (* $p < 0.05$, ** $p < 0.01$ and *** $p < 0.001$)
